# Hidden in Plain Sight. How Ks histogram dynamics can reveal and obscure ancient whole genome duplications

**DOI:** 10.1101/2025.11.19.689290

**Authors:** Tamsen Dunn, Arun Sethuraman

## Abstract

Polyploidy is a significant force in angiosperm evolution, of great interest to evolutionary biologists and crop scientists. The mode of origin of a polyploid lineage is a continuum, ranging from whole genome duplication (WGD) from within the same species (autopolyploidy), to a WGD derived from hybridization between two (or more) distinct species (allopolyploidy). The polyploid mode of origin impacts many aspects of genome evolution, including patterns of chromosomal inheritance, homeologous exchange rate, and post-WGD diploidization. Here, we develop a novel polyploid genome simulation engine DemographiKS to demonstrate that the Ks histogram, one of the most broadly utilized methods for inferring WGD events, is highly sensitive to evolutionary parameters relating to the mode of origin, as well as demographic parameters such as migration events, population bottlenecks and expansions. Our results show that that the location of the Ks peak may correspond to the time of parental divergence (for allopolyploids); might be hidden at Ks=0 (for autopolyploids under a population contraction); may correspond to the mean time of coalescence of the diploid ancestor (for autopolyploids under a population expansion), or the time of most recent migration between the parental species (for allopolyploids). We also fit simulated Ks histograms to polyploid genomes from four well-studied plant lineages on the polyploid continuum (*Coffea arabica*, *Zea mays*, *Populus trichocarpa*, and *Saccharum spontaneum*), demonstrating that WGD simulations may be used to corroborate inferred demographies. In summary, we demonstrate that Ks histograms are information-rich, computationally tractable, and can be utilized to corroborate detailed evolutionary histories inferred by other methods.

## Introduction

One of the great success stories of evolution is the explosive diversification and dominance of the angiosperms. Many explanations have been given for angiosperms’ success (1–5). One plausible explanation is that the key to angiosperm success is the numerous rounds of whole genome duplication (WGD) that have occurred throughout their evolution (6–13). Additionally, since all crop plants are polyploid derived species, untangling how WGD may contribute to their fitness and phenotypic novelty is of major interest to agricultural scientists seeking to improve yield and stress tolerance (14–21).

When a genome duplicates, the additional genome(s) may be sourced from within the same species (autopolyploidy) or from a closely related species (allopolyploidy) (22–25). While strict categorization of “auto” or “allo” might not reflect the reality of biologically ‘messy’ speciation processes, these binary designations are useful because they allow us to identify and contrast the effects of higher and lower levels of diversity between ancestral genomes (25–29). Real polyploids exist in a continuum of multiple parameters (27, 28, 30–37). To that end, we present the novel polyploid simulator DemographiKS which allows polyploid evolution to be modeled as a ‘messy’ continuum, where the mode of origin, level of differentiation between genomes, levels of homologous and homeologous exchange, and proportion and timing of gene flow by secondary contact may all be parameterized by the user (software architecture, Supp. Fig. S1, underlying model Supp. Fig. S2).

Historically, allopolyploidy has been considered more common than autopolyploidy, a view point advocated by G. Ledyard Stebbins and his contemporaries (38–43). However, recent work illuminating the complexity and subtlety of the determining the mode of paleopolyploid speciation only underscores that we still do not have a reliable estimation of the relative proportion of speciation by allopolyploidy or autopolyploidy across the plant tree of life (29, 32, 41, 44–47), and several works (including this one) suggest that autopolyploidy might be under-detected (32, 34, 45).

While the effects of auto and allopolyploidy have many similarities (for example, both increase overall organ size, increase in the potential number of mutational targets and may reduce the power of selection, (23, 48, 49), there are many differences, specifically with respect to the process of diploidization post-WGD (28, 30, 50). Diploidization is the gradual process by which a polyploid lineage returns to diploidy. This critical process is characterized by intense genomic reorganization, where old genes are shed, and new genes and phenotypes may be formed at greater rates (2, 3, 8, 9, 12, 36, 48, 51, 52, 52–60).

The differences in diploidization between auto and allopolyploids largely has to do with how well duplicate genomes ‘play together’. For autopolyploids, there are no distinguishable subgenomes. This poses a significant problem for reproduction, since chromosomes have multiple potential homologous partners (polysomic inheritance), instead of a single partner during meiosis (disomic inheritance). Polysomic inheritance can result in multivalent chromosomes forming chains and rings, instead of the orderly pairs formed by bivalent pairing. The impact of polysomic inheritance on fertility varies by lineage, and research shows that autopolyploids generally return to fertility once gametogenic stability is achieved. Thus, for many autopolyploids, removing redundancy and achieving differentiation between homologs is key to diploidization. (34, 57, 61–72).

In contrast, the duplicate genomes of allopolyploids may already be separated by several million years of evolution. Allopolyploids with a full diploid set of chromosomes from distant relatives typically have few issues with fertility (23). For these allopolyploids with even numbered chromosomes inheritance is disomic, though the rates of homeologous exchange across subgenomes may vary (35, 50, 73–78). Polyploids with odd numbers of chromosomes face a unique crisis where chaotic chromosomal segregation seems inevitable. However, research has shown that even for triploids, different lineages display varying levels of fertility and can return to diploidy (79–82).

With respect to fitness, many studies have shown that allopolyploidy provides an injection of hybrid vigor in the form of heterosis (12, 83–85). However, this benefit may be countered by a range of adverse effects due to the long separation endured by the parental genomes before hybridization. The neopolyploid may experience increased transcriptomic shock, transposable element release, cytonuclear incompatibilities and stoichiometric imbalances in signaling and regulatory processes (82, 86–90). Thus, for an allopolyploid, a critical aspect of diploidization is a return to orderly genome management (58, 90–94).

Modern techniques for detecting WGD events generally fall into three categories: synteny based methods, phylogenetic methods, and Ks-based methods. Syntenic methods identify conserved ordering in blocks of genes between putative subgenomes and between related species (95–102). Phylogenetic methods use the comparative analysis of gene trees to bind gene duplication events to specific regions on a phylogenetic tree, then seek to reconcile the gene duplication events to a common species-level topology (28, 103–113).

Ks-based methods, first demonstrated by Blanc and Wolfe in 2004, are arguably the simplest and most widely used. However, no Ks-based methods exist yet that can accurately incorporate the complexity of the polyploidy continuum (114–118). Here we develop the novel forward-time polyploid continuum simulator DemographiKS to demonstrate that the Ks histogram is sensitive to many demographic and evolutionary parameters of interest, including mode of origin and homeologous exchange rate. In **part 1** of the results section, we validate DemographiKS against theory and prior simulations. We do this by showing that the DemographiKS Ks peak placement and histogram shape match both theoretical predictions under a simplified evolutionary model, and also match predictions from the previously-published SpecKS (Dunn and Sethuraman 2024), under similar constraints. In **part 2**, we move beyond simplifying assumptions, to explore the effects of critical genetic and evolutionary processes not included in prior work. We explore the effects of (a) recombination, (b) migration, (c) homeologous exchange, and (d) population bottlenecks and expansions. Additionally, in each subsection, we compare the simulated results from DemographiKS with theoretical expectations. In **part 3**, we compare and contrast DemographiKS results with empirically observed Ks histograms, for four different ancestral polyploids: *Coffea arabica*, *Zea mays*, *Populus trichocarpa*, and *Saccharum spontaneum*. In conclusion, we describe future work needed to close the gap between DemographiKS results and biological observations.

## Results

### Part 1: Validation of DemographiKS against prior work and theory

We compare DemographiKS results against theory and prior computational studies via demonstrating that the DemographiKS Ks peak placement and histogram shape matches both theoretical predictions given a simplified evolutionary model (propagation of the Kingman coalescent through time), and predictions from the previously - published SpecKS, under similar evolutionary constraints. In row one of Fig. 1, we show how the tail of the Ks distribution lengthens as a response to higher ancestral effective population size (N_a_), where the mean of the exponential distribution matches the theoretical prediction of 2N_a_. In row two of Fig. 1, we show, as more time passes, the structure Ks histogram erodes as the signal attenuates over time. Over larger time scales, the mean of the Ks distribution is given by the time since parental divergence, in Ks space, which also matches theoretical predictions given by (Dunn and Sethuraman 2024). Concordance between theory and simulation in response to varying effective population size and time since parental divergence has been established previously for SpecKS. However, our experiment additionally demonstrates that DemographiK is also concordant with both SpecKS and theory, for three orders of magnitude for variations in both population size (*N_a_* =10, 100, 500, 100) and time since parental divergence (*T_DIV_* = 10000, 100000, 50000, 100000). Simulation input parameters are given in Supp. Tables S1-S4.

**Figure 1.**
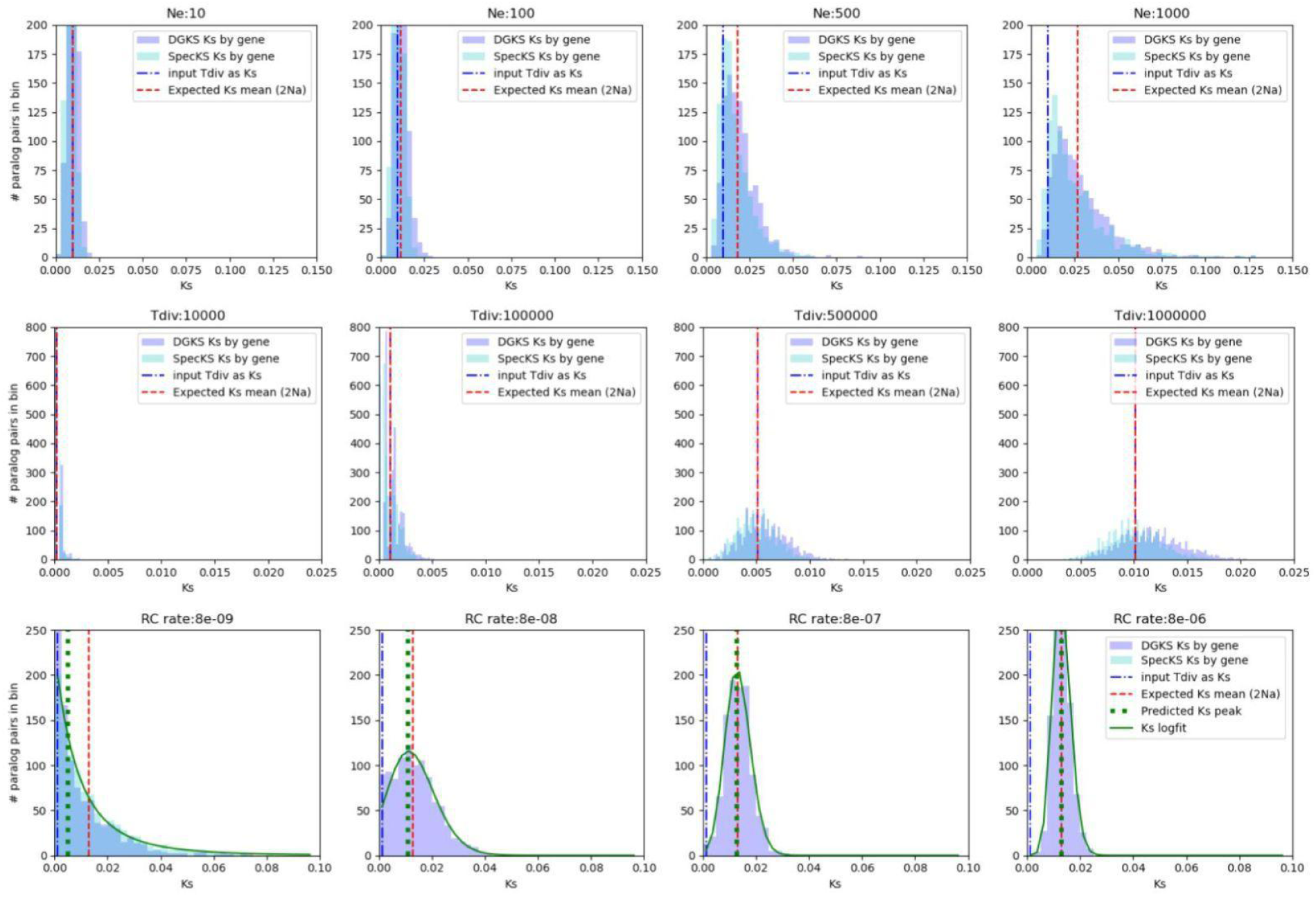
Demographiks, specks and theoretical expectations are in agreement, for variation in population sizes and time scales, and when recombination rate is low. Row 1: DemographiKS, SpecKS and theoretical expectations are in agreement for variations in N_e_. All demonstrate that increases in N_e_ increase the skew of the Ks histogram. Here we show simulated Ks distributions 1000 generations after WGD, for N_e_ ranging from 10 to 1000. Row 2: DemographiKS, SpecKS and theoretical expectations are in agreement for variations in time since parental divergence (T_DIV_). All demonstrate that increases in T_DIV_ shift the Ks histogram to the right. The simulated Ks distributions are for N_e_=1000, with time since WGD/Tdiv ranging from 10K to 1M generations ago. Row 3: DemographiKS, SpecKS and theoretical expectations are in agreement for variations in recombination rate. All demonstrate that as recombination rate increases, the Ks distribution loses skew, changing shape from exponential, to lognormal, to Gaussian. Here we show simulated Ks distributions for simulations with N_e_=10000, for DemographiKS recombination rates ranging from 8×10^-9^ to 8×10^-6^ per base pair per generation. Note that on the leftmost panel, we use SpecKS (recombination rate=0) along with DemographiKS recombination rate (8×10^-9^). SpecKS does not appear in any other panels on row 3. This is because SpecKS does not model recombination (i.e. recombination rate is fixed at 0). DemographiKS utilizes a non-zero model of recombination required to break up linkage disequilibrium. For simplicity, time of parental divergence across all figures is set to time of WGD (Δ*T* = 0), and there is no homeologous exchange between subgenomes. Predictions for the mean coalescent given by the Kingman coalescent are given by the red dashed line. Model predictions of the leading edge of the Ks distribution, based on parental divergence time, is shown by the blue line. DemographiKS results are shown in purple. SpecKS results are shown in cyan. Predictions of the Ks histogram peak including recombination are given by the green dashed line, and the lognormal fit in solid green.

### Part 2: The effects of recombination, migration, homeologous exchange and changes in population size on the Ks histogram

Here we explore the effects of critical genetic and evolutionary processes of (a) recombination, (b) migration, (c) homeologous exchange, and (d) population bottlenecks, and expansions on the shape and peak placement in Ks histograms. Additionally, we compare the simulated results from DemographiKS with theoretical expectations.

#### Recombination

Since recombination is iterative resampling of chromosome segments from a population, we theorized that, by virtue of the central limit theorem, the original Ks distribution would tend towards a Gaussian distribution over time, as more recombination events reshuffled the polyploid genome (Supp. Fig. S3 and S4). This predicted reshaping was confirmed by DemographiKS. As expected, our results (Fig. 1, row 3) confirm the expected relationship between recombination rate and histogram peak location: when the recombination rate is zero, the Ks histogram is determined by the ancestral Kingman coalescent distribution, peaking at the time of parental divergence. As the recombination rate increases, the distribution tends towards a Gaussian distribution, with the histogram peaking at the mean coalescent time. In this figure, both SpecKS and DemographiKS matched theoretical predictions in their respective domains. Simulation input parameters are given in Supp. Tables S5 and S6.

#### Migration

We anticipated that when migration occurs between the ancestral diploid parental species of an allopolyploid, after initial parental divergence but before the genomes came back together due to WGD, this process would shift the Ks peak leftwards, closer to the time of WGD. This is because when introgression between the parental species occurs, some orthologs between the parental species might have common ancestry tracing back to the time of migration, rather than the earlier speciation event between the parental species. After whole genome duplication has occurred and the orthologs ‘become’ paralogs (ohnologs), and this would translate into some ohnologous pairs tracing their common ancestor back to the time of migration, rather than the earlier speciation event between the diploid parental species (Supp. Fig. S5).

To test the sensitivity of the Ks histogram to migration arising from secondary contact as described above, we set up two experiments, investigating (1) stretches of secondary contact spanning 5 generations, and (2) stretches of secondary contact spanning 100 generations, both starting at the halfway point between the parental divergence time and the present day. Our findings indicate that even small amounts of migration (1% at 5 generations post divergence) provide detectable shifts in the Ks histogram. As expected, the greatest amount of migration caused the strongest perturbation of the Ks histogram peak. Interestingly, for protracted secondary contact (100 generations post divergence) the effects of a 1% migration rate are much the same as a 50% migration rate (Supp. Fig. S56 and S7, with parameters given in Supp. Table S7).

To test the sensitivity of the Ks histogram to gradual speciation modeled as low-level ongoing migration after parental divergence (Isolation with Migration) we set up a third experiment. Here we simulated continual migration of 0.1%, 1.0% and 10%, lasting for 50K, 100K, 250K, 1.0 M generations. Results from our gradual speciation with gene flow experiments indicate that long term migration, even at low rates, are easily detectable if it lasts for a long enough period of time (Fig. 2, parameters given in Supp. Table S7). However, we also see that when migration concludes soon after parental divergence, perturbations to the Ks histogram are difficult to detect, even with a strong migration rate (10%). This latter finding is expected because if migration concludes relatively quickly, the new Ks peak will not be far from the original peak, so perturbations will be difficult to detect. (For example, if the width of the Ks peak itself is the equivalent of 500K generations, and migration concludes after 5K generations, the peak would not move left more than 5/500, or 1% of the original Ks peak width.

**Figure 2.**
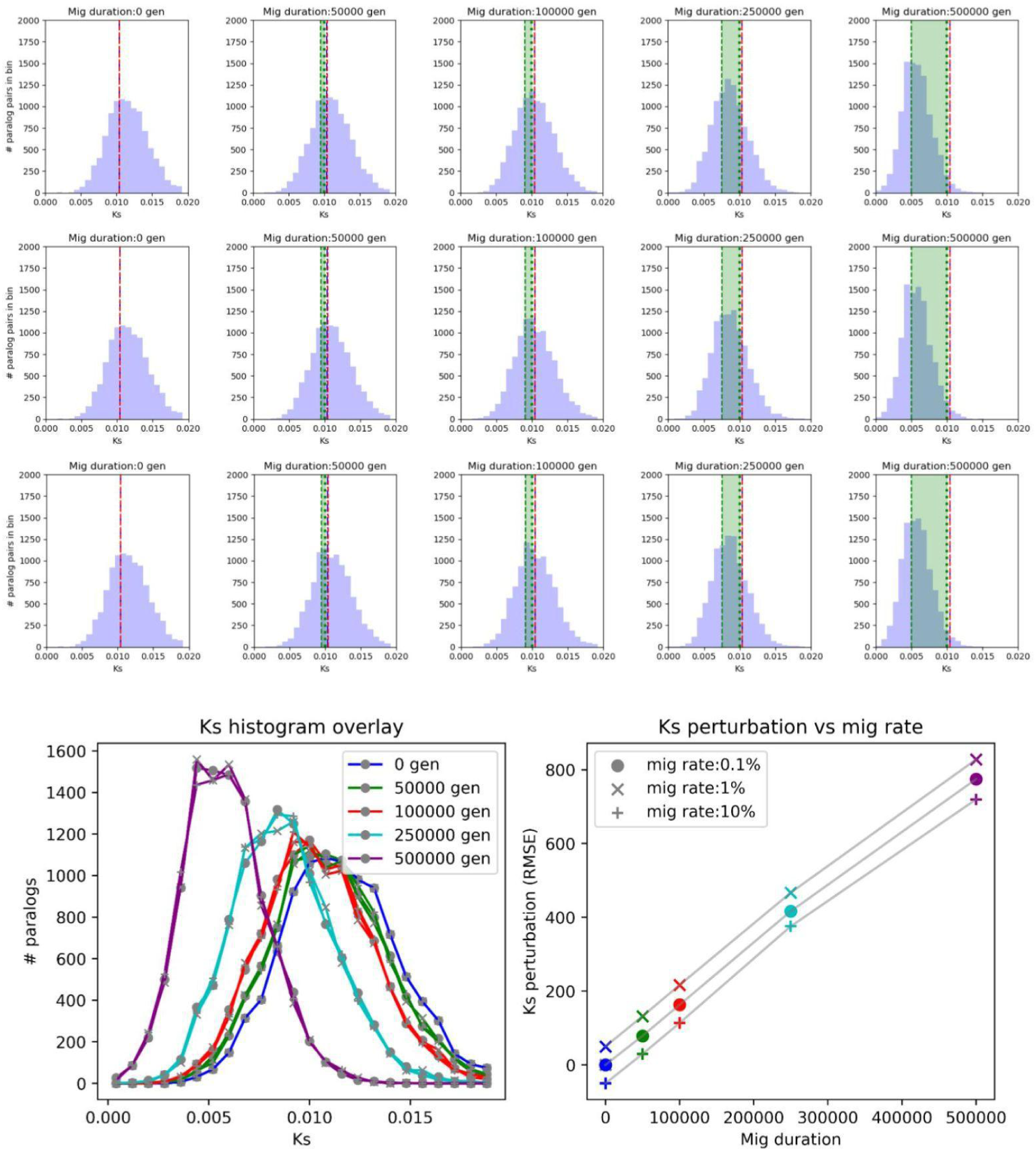
Prolonged gradual speciation even at low percentages, can significantly shift the Ks histogram. Rows 1-3: Ks histograms for migration of 0.1,1.0 and 10%, lasting for 50K, 100K, 250K and 0.5 million generations. The red dashed line indicates the predicted distribution center of mass, without migration. The green dashed line indicates the time of migration, in Ks-space. The duration of migration is overlaid in green shading. Bottom left, overlay of Ks histograms. Line colors correspond to the duration of migration and marker style corresponds to migration rate. Bottom right: Perturbations between the null hypothesis Ks histogram (no migration) and the given migration rate Ks histogram, measured as RMSE. Marker colors correspond to the duration of migration. Marker style corresponds to migration rate. The lines on the bottom right figures are artificially displaced, and would otherwise lie directly on top of each other.

#### Homeologous exchange

The effects of homeologous exchange on the Ks histogram are not well understood. In general, one might expect homeologous exchange to be more common in autopolyploids than allopolyploids, due to tetrasomic inheritance in the former and disomic inheritance in the latter. Thus, under the rules of tetrasomic inheritance, one would expect the mean coalescent time between ohnologous pairs of genes to be 4N_e_, analogous to how one expects a mean coalescent time of 2N_e_ under disomic inheritance (119). Therefore we predicted that for idealized autopolyploids under high levels of homeologous exchange, the simulated Ks histogram should match the expected Kingman coalescent under tetrasomic inheritance, with peak at Ks=0 and mean of 4N_e_, where N_e_ is the effective population size of the present polyploid population. Our expectations for (idealized) autopolyploids and allopolyploids are very different. For allopolyploids, the Ks histogram peak represents the time of ancestral divergence (see T_DIV_ from (28, 114).

Indeed, when DemographiKS is used to simulate the Ks histograms of polyploids with high levels of homeologous exchange, the resulting histogram conformed to a Kingman coalescent distribution with peak at Ks=0 and mean of 4N_e_, confirming our theoretical expectations that the peak would not occur at the time of ancestral divergence. Here we tested polyploid populations of sizes 50, 100, 500 and 1000 (Fig. 3, top). We did not see a secondary peak corresponding to the time of whole genome duplication or the time of parental divergence (Fig. 3, top). Unlike the allopolyploid Ks histogram (Fig. 3, bottom, purple) the autopolyploid Ks histogram showed no dependency on time of whole genome duplication (Fig. 3, bottom, green), even after one million generations simulated. Simulation parameters when the population size is varied are given in Supp. Table S8, and when time since parental divergence/whole genome duplication is varied is given in supplemental Supp. Table S9.

**Figure 3.**
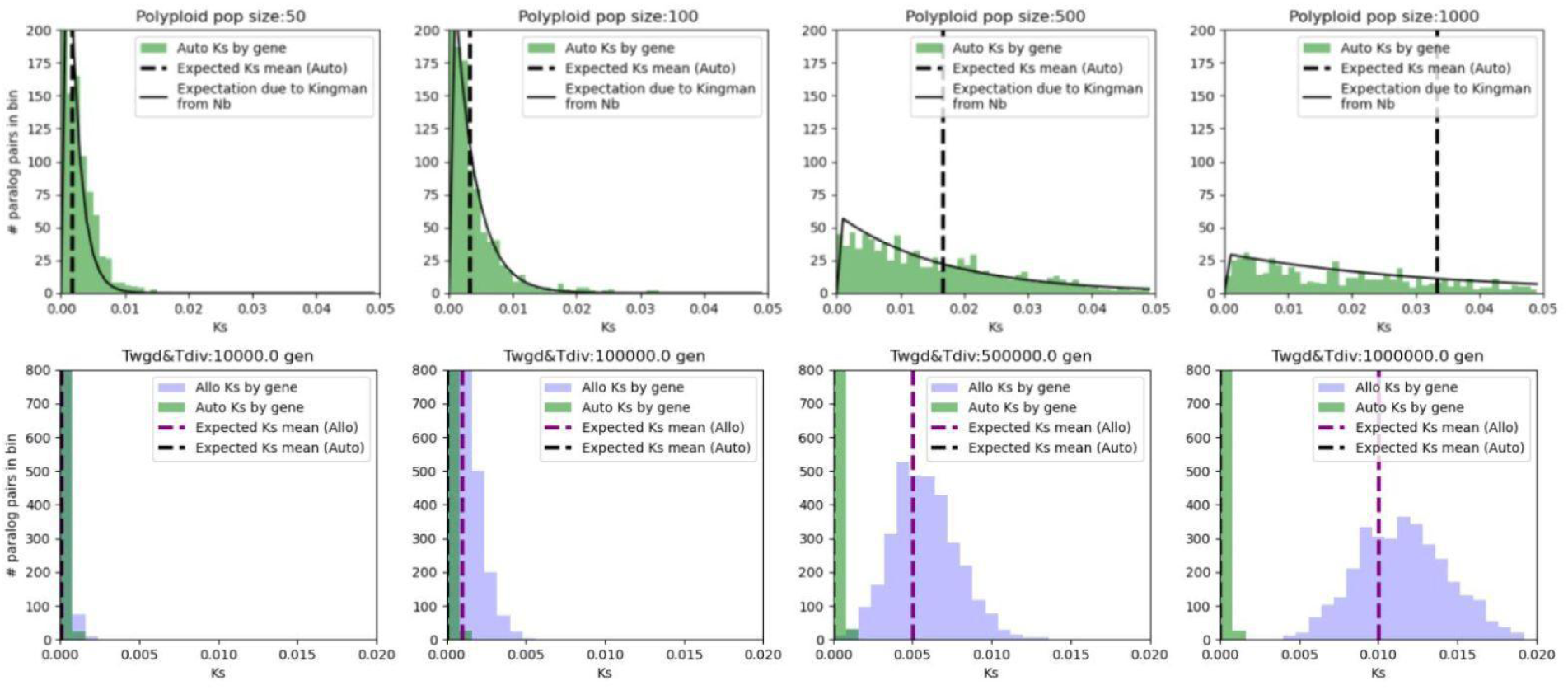
Top: The simulated Ks distribution for polyploids with free homoeologous exchange matches the expected Kingman coalescent, across varying polyploid population sizes. The following Ks histograms are for polyploid populations of sizes 50, 100, 500 and 1000. In all simulations, there is no peak at the Ks value corresponding to the time of parental divergence, nor at the Ks value corresponding to the time of whole genome duplication. The solid black line represents the expected Ks histogram due to the Kingman coalescent. The dashed line represents the predicted mean coalescence. **Bottom: As expected, the simulated Ks distribution for polyploids with free homoeologous exchange (green) has no time dependencies.** In contrast, the Ks distribution for polyploids without homoeologous (purple) exchange moves right along the x-axis as time since parental divergence increases. Here, the input time since WGD (T_WGD_) was 10K, 100K, 500K, and 1M generations ago. For the allopolyploid case, T_DIV_ = T_WGD_, hence (Δ*T* = 0), and for the autopolyploid case, time since parental divergence was set to FALSE.

**Figure 4.**
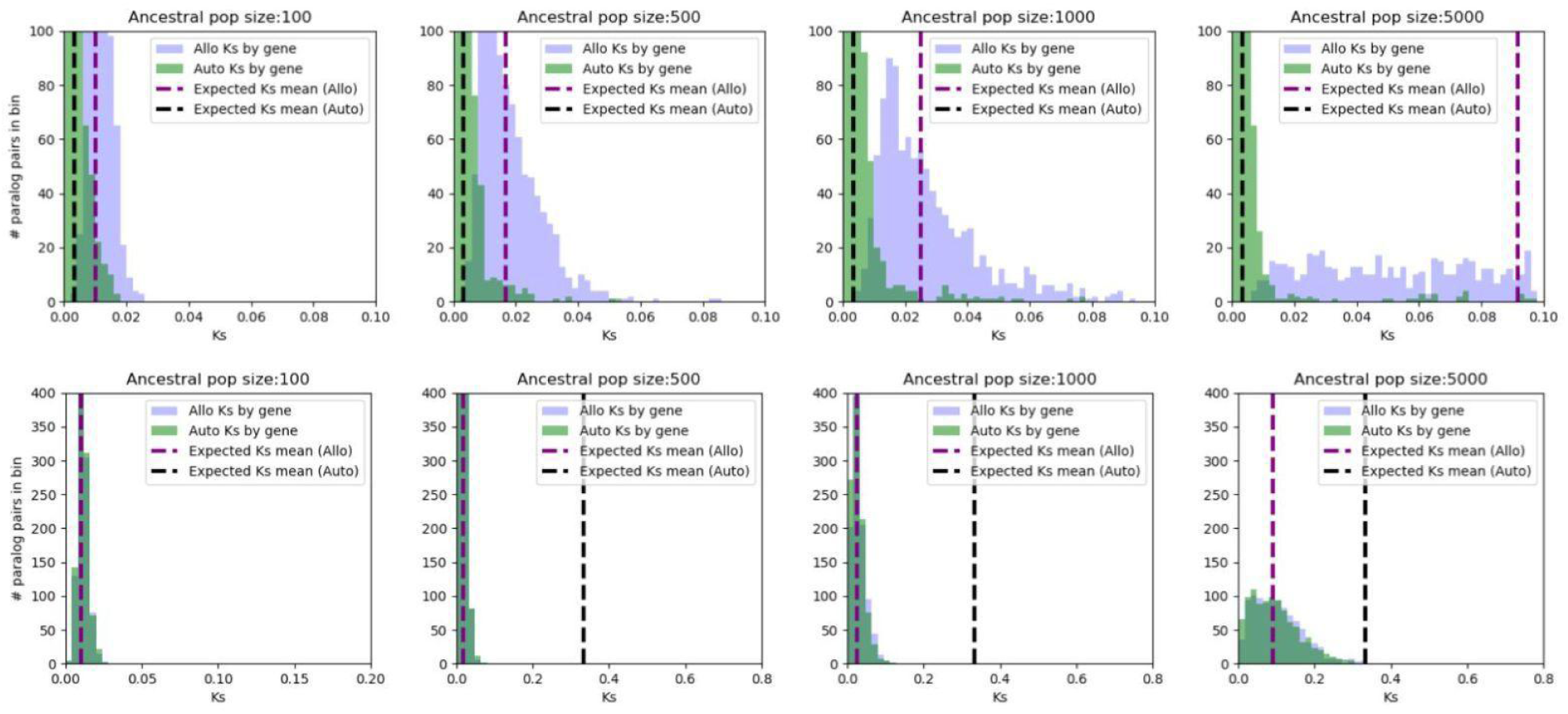
Simulated Ks histograms for auto and allopolyploids can look very different or extremely similar, depending on whether the population is under expansion or contraction. **Top: When N_b_ < N_a_, the allo and autopolyploid Ks histograms look very different, as expected.** For the allopolyploid, the Ks centroid (the mean coalescent) follows predictions based on N_a_, (purple dashed line), while for the autopolyploid the Ks centroid follows predictions based on N_b_ (black dashed line to the left). Here N_b_ is held at a constant 100, and N_a_ varies from (100, 500, 1000, 5000). **Bottom: However, when N_b_ >> N_a_, the auto and allopolyploid histograms look very alike, with both showing a strong response to ancestral population size.** Here for the allopolyploid, the Ks centroid again follows predictions based on N_a_ (purple dashed line), but what is surprising is that the autopolyploid also follows predictions based on N_a_. This is because the expected mean coalescence based on N_b_ is further back in time (black dashed line to the right), thus is superseded by the more recent coalescence of the ancestral population. Here N_b_ is held at a constant 10K, and N_a_ varies from (100, 500, 1000, 5000) individuals.

#### Population bottlenecks and expansions

It has been previously demonstrated that for allopolyploids, the Ks peak would correspond to the time of parental divergence, and that smaller population sizes would shorten the tail of the Ks distribution and larger population sizes would lengthen the tail (114). Those results were in line with theoretical expectations, which predict that the effects of population size on the Kingman coalescent would directly translate into similar effects on the Ks histogram, in Ks space. However, there are two significant limitations with prior work: firstly, homeologous exchange was not taken into account, and secondly, population size was considered constant throughout the time period of the simulation. We expected that when ancestral population size and polyploid population size were decoupled, as in DemographiKS, ancestral population size rather than polyploid population size would determine the shape of the Ks histogram when there was no homeologous exchange (in agreement with previous work (37), and Fig. 1). We additionally predicted that polyploid population size rather than ancestral population size would determine the shape of the Ks histogram when homeologous exchange occurred freely (in agreement with Fig. 3). Thus, the genomic signatures of bottlenecks and expansions might look very different for auto and allopolyploids. Our predictions and results are summarized in (Supp. Table S10).

As expected, our results showed that when ancestral and polyploid population sizes are allowed to vary independently, allopolyploid Ks histograms shapes are exclusively determined by ancestral population size (Supp. Table S10 and Supp. Fig. S8, row 1 (constant N_b_=100, N_a_ varies), row 2 (constant N_b_=10K, N_a_ varies). Polyploid population size has very little effect (Supp. Table S10 and Supp. Fig. S8, row 3 (constant N_a_ =100, N_b_ varies), row 4 (constant N_a_ =10K, N_b_ varies)). This is consistent with prior work (114) and our comparisons with SpecKS (Fig. 1).

DemographiKS additionally confirmed that when homeologous exchange is freely allowed, such that there are no maintained subgenomes, the shape of the Ks histogram is dependent on the current polyploid population size (Supp. Table S10 and Supp. Fig. S9, row 3 (constant N_a_ =100, N_b_ varies), and row 4 (constant N_a_ =10K, N_b_ varies). We also observed that, as expected, for relatively small polyploid population size (N_b)_) (Supp. Table S10 and Supp. Fig. S9, row 1 (constant N_b_=100, N_a_ varies)), variation in ancestral population size had little effect. This is also consistent with expectations based on models of tetrasomic inheritance, shown in Fig. 3.

Lastly, for situations where homeologous exchange occurs and N_b_ is large relative to N_a_, we expected that there might be coupling between the coalescence such that N_a_ would still affect N_a_ (Supp. Table S10, marked with “??”). We wanted to explore how the Ks histogram would react when the mean ancestral coalescence time occurred more recently than the expected mean polyploid population coalescence time. For example, if the polyploid population size is 5,000, the expected mean coalescent time is 4N_e_, so we would expect randomly selected ohnologs to coalesce 20,000 generations before the present. However, if the ancestral population was quite small (say, 100), then any two randomly selected gene pairs from the ancestral population would be expected to coalesce only 200 generations back. This sets up a contradiction. Because no matter whether the polyploid is auto or allopolyploid, the mean coalescence cannot go back further in time than 200 generations back from the original speciation time. Future events like population expansions cannot push a coalescent back in time further than the ancestral population supports.

Thus we expected that under the paradigm of a rapid, recent population expansion, the expected tail of an autoploid Ks histogram would be truncated due to the small ancestral population size, such that it could never expand further back in time past the predicted allopolyploid Ks histogram for the same ancestral populations size. Mathematically, we expected that if 4N_b_ > T_spec_ + 2N_a_, the autopolyploid Ks histogram would be cut short, to overlap with the corresponding allopolypoid Ks histogram.

To test this, we used DemographiKS to run a series of experiments where the current polyploid population size was held constant (at either 100 or 10,000) and the ancestral population size was varied between 100, 500, 1,000 and 5,000. As expected, when the polyploid population size was very small (100), the auto and allopolyploid Ks histograms were very different, with only the allopolyploid showing a strong response to ancestral population size (Fig. 4, top). However, when the polyploid population size was very large (N_a_ = 10,000), the auto and allopolyploid Ks histograms overlapped almost perfectly, with both showing a strong response to ancestral population size (Fig. 4, bottom, and Supp. Fig. S9, row 2 (constant N_b_=10K, N_a_ varies)). However, when the polyploid population size was very small (N_b_ =100), the autopolyploid Ks histogram remains peaked near Ks=0 and the allopolyploid Ks peaks near the time of parental divergence, with a shape corresponding to the ancestral population size (Fig. 4, bottom, and Supp. Fig. S8, row1 (constant N_b_=100, N_a_ varies)). Thus we report novel coupling effects between ancestral and modern population sizes for autopolyploids under population expansion. (Parameters given in Supp. Table S11).

### Part 3: A comparison between DemographiKS results with empirical Ks histograms

We stress that the concordance between our idealized expectations for canonical auto and allopolyploids given in part 2 will not map directly to concordance with expected results for real polyploids. Real polyploids exist as points in a continuum of allo and autopolyploid traits, with segments of the genome operating differentially across time and across the genome. One set of homeologous chromosomes might have a different rate of homeologous exchange from another set, due to the different levels of differentiation between the chromosomes in the sets. Furthermore, those rates of exchange may change over time as diploidization affects differentiation, since some sets of chromosomes may initially have tetrasomic inheritance, but over time and due to differentiation, move to disomic inheritance. Thus in practice, empirical Ks histograms may show signatures of a mixture of processes, on different timescales. DemographiKS also does not currently include gene pairs maintained by selection, which has been theorized to account for the long flat tail seen in empirical Ks distributions (115, 120).

Thus we expected that allowing variation in patterns of homeologous exchange and including effects from gene pairs maintained over time by selection might capture missing features of the DemographiKS Ks histogram. To test this, we compared (a) default DemographiKS Ks histograms with no homeologous exchange, (b) Ks histograms from a combination DemographiKS results, where some portions of the genome had no homeologous exchange and in others homeologous exchange was enabled, and (c) additionally including a percentage of paralogs to be born from a random birth-death model, some of which would be never be shed (115, 121).

In order to test whether DemographiKS results would deviate from empirical observations according to our expectations, we selected four ancestral tetraploids: *Coffea arabica, Zea mays, Populus trichocarpa,* and *Saccharum spontaneum*. These ancestral polyploids are well-studied, and in our simulations we used previously-inferred values as input parameters wherever possible (Supp. Tables S12-S15). In our first set of tests, we used a single run of DemographiKS to simulate the Ks histogram for each species. Without including homeologous exchange, we were able to achieve RMSE fits of (1.84, 2.23, 1.94, 2.02) for *Coffea arabica, Zea mays, Populus trichocarpa,* and *Saccharum spontaneum*, respectively. *Coffea arabica* had the best fit (RMSE=1.84), which was expected because *C. arabica* is a recent allopolyploid (122, 123), so there has been little time for homeologous exchange and diploidization. Conversely, *Z. mays* and *S. spontaneum*, both ancient polyploids with complex histories (124–135) proved the most difficult to model (RMSE=2.23 and 2.02, respectively). (Fig. 5, first column).

**Figure 5.**
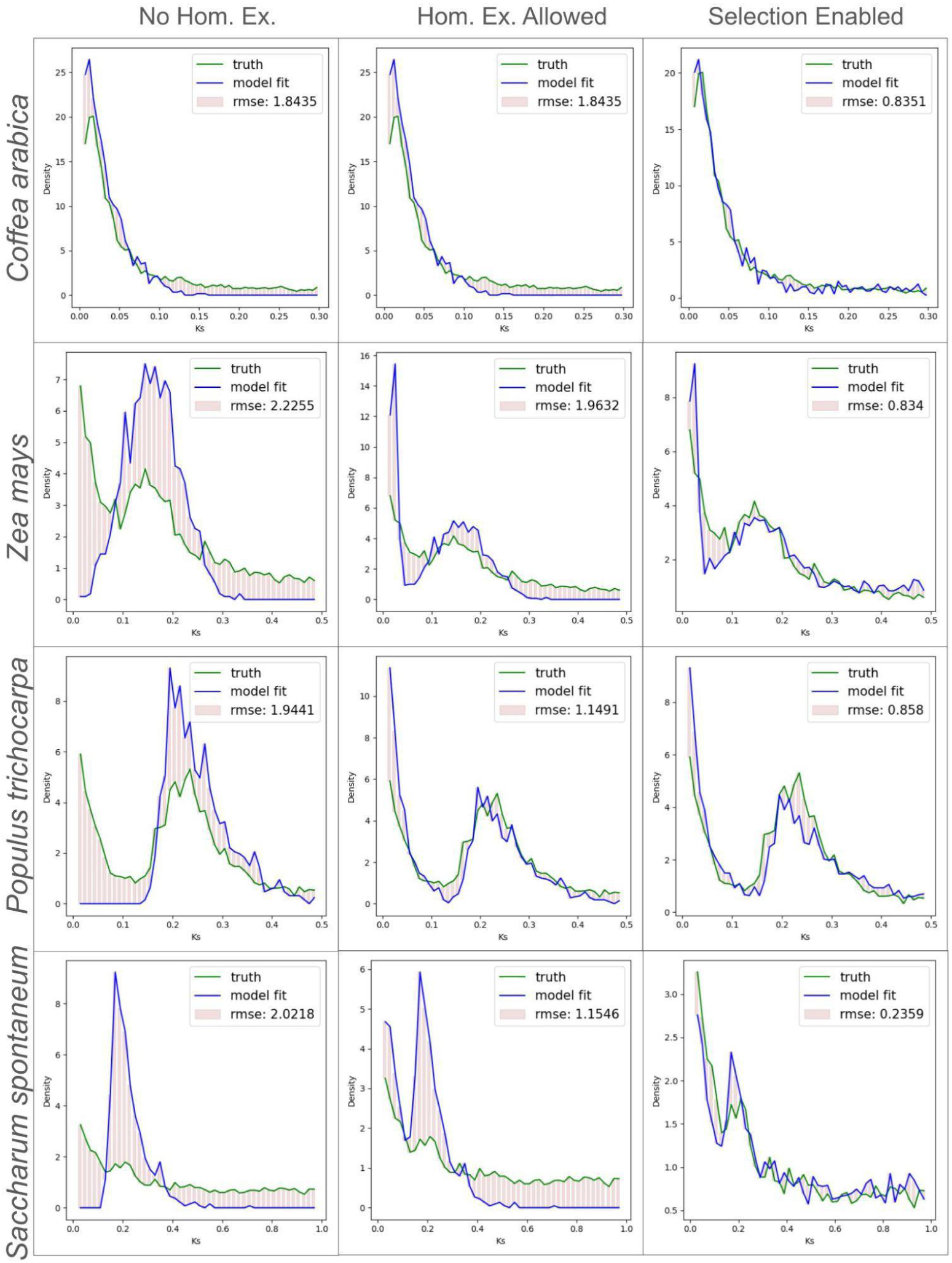
Incorporating homeologous exchange and selection into our model provides a better fit to empirical results than a single, basic DemographiKS run. Here we give the empirical Ks distributions in green and our simulated results in blue. Fit parameters are given in Supp. Tables S16-S19 The x-axis represents Ks and the y-axis represents density. The pink rectangles represent the root mean squared error (RMSE).

In our second set of tests, for *Z. mays*, *P. trichocarpa* and *S. spontaneum*, we combined results from two runs of DemographiKS, to model the effects of different subsets of the genome having different homeologous exchange rates, again using previously-inferred values as input parameters wherever possible. By including homeologous exchange, we were able to achieve RMSE fits of (1.96, 1.15, and 1.15) for *Z. mays*, *P. trichocarpa*, and *S. spontaneum*, respectively, with error rates dropping on average by about 30%. (Fig. 5, second column).

Lastly, we expected that DemographiKS fits could be further improved by allowing some portion of paralogous genes to be maintained by selection, rather than shed over time. Thus in our third set of tests, we included a set of Ks values corresponding to maintained paralogs which arise due to selection favoring and maintaining a fraction of continuously-arising small scale duplications. This is well described in (115), and we describe our mathematical model in the supplemental methods section. As expected, adding maintained paralogs allowed the right-hand tail of the simulated Ks histogram to asymptotically reach equilibrium over deep time (as observed in empirical datasets), rather than trail off to zero (as we saw in our first and second test sets.) By including selection in this manner, we were able to achieve RMSE fits of (0.86, 0.83, 0.86, 0.24) for *C. arabica, Z. mays, P.trichocarpa,* and *S. spontaneum*, respectively. This decreased the error on average by an additional 50%. (Figure 5, third column). All DemographiKS parameters for these simulations are given in Supp. Tables S16-S19).

## Discussion

Here, we present the polyploid continuum genome evolution simulator DemographiKS. DemographiKS is the first Ks simulator that can effectively marry population, genome, and molecular evolutionary dynamics in a polyploid simulation engine. The provenance of such a Ks simulator is timely. Observed Ks histograms are now abundant and relatively easy to generate (118, 136, 137). DemographiKS can be used to derive an estimated Ks histogram, and comparisons between the observed Ks histogram and estimated Ks histogram can then be made to corroborate or refute demographic inferences, and assess how sensitive the estimated Ks histogram might be to uncertainty surrounding input parameters. For example, a comparison between the observed Ks histogram and a range of estimated Ks histograms might be used to examine whether a migration event might have occurred, and for how long, or to test how frequent historical homeologous exchange might have been between the subgenomes.

In this paper we have explored the effects of (a) recombination, (b) migration, (c) homeologous exchange, and (d) population bottlenecks and expansions. Additionally, in each subsection, we compare the simulated results from DemographiKS with theoretical expectations. We have demonstrated that the Ks distribution represents a powerful and complex distribution sensitive to many aspects of ancient demography. We have shown how Ks histogram dynamics both reveal and obscure ancient whole genome duplication events. For example, the Ks histogram peak may correspond to the time of parental divergence (for allopolyploids); might be hidden at Ks=0 (for autopolyploids under a population contraction); may correspond to the mean time of coalescence of the diploid ancestor (for autopolyploids under a population expansion), or the time of most recent migration between the parental species (for allopolyploids).

We also fit simulated Ks histograms to empirical results from four well-studied plant lineages (*Coffea arabica*, *Zea mays*, *Populus trichocarpa*, and *Saccharum spontaneum,* Fig. 5), demonstrating that simulations may be used to corroborate inferred demographies. In summary, we demonstrate that Ks histograms are information-rich, computationally tractable, and can be utilized to corroborate detailed evolutionary histories inferred by other methods. Used with appropriate caution, the Ks histogram may serve as a companion to other methods such as the site frequency spectrum to corroborate demographic inference in polyploid species.

In future work, we aim to develop a formal and scalable framework for determining the optimal parameters which allow DemographiKS to most closely model the empirical Ks histogram for a given lineage. Since both the estimated Ks histogram and observed Ks histogram are relatively easily calculated, one could write an objective function based on the agreement between the observed and estimated Ks histograms, and set up an input-parameter-estimation loop, leveraging existing numerical optimization approaches. Additionally, DemographiKS may be extended to include other biological processes (such as population structure, selection or multiple rounds of WGD).

Including more complexity in our models would increase computational time. However, we anticipate that as theoretical advances are made to Ks histogram estimation, there should be less reliance on individual-based simulation, which would reduce computational load.

### Caveats

Here we give a brief overview of caveats to consider when using DemographiKS. A more detailed exposition is included in the ‘discussion section of the supplemental materials.

A significant accumulation of synonymous mutations are needed to resolve the shape of distant effects on the ancient coalescent. If the ancestral coalescent is too shallow or the mutation rate too small, the number of mutations collected during ancestral speciation may be too small to affect the final Ks distribution. We also acknowledge DemographiKS is slow compared to SpecKS. To save computational time, small populations and short time scales were used in this paper wherever possible. In the next generation of Ks simulators, we anticipate that the estimation of Ks distributions might rely more on theory and less on individual-based simulation, and thus be faster to compute. An additional limitation of the DemographiKS model is that we presume that the rates of recombination, mutation and homoeologous exchange are constant over time and spatially along a chromosome. We also assume no population structure. The effects of the rates of various parameters changing over time will surely integrate over long timescales (30, 105, 138). It is also possible that different theoretical evolutionary histories might create the same Ks histogram. We also noted that in our empirical fits, we frequently had to use a lower recombination rate than was suggested in the literature, else the histograms would tend to Gaussians faster than expected.

One explanation for this might be that our model of recombination did not take into account population structure, specifically isolation by distance. Since in real populations, recombination would be happening between similar genomes, rather than between random genomes selected from within the entire population, we might expect that in our simulations, recombination had a stronger effect than would be observed in nature. Future work will iteratively need to incorporate additional complexity into a new generation of polyploid simulation tools. In reality, particularly within a polyploid genome, we expect that many of these parameters will change over time, due to diploidization and general molecular evolutionary effects.

## Methods

### Software implementation

DemographiKS is implemented as a pipeline application in python3. It wraps three major components: SLiM, msprime, and PAML, as shown in Supp. Fig. 1 (139–141). DemographiKS takes as input an XML configuration file listing the simulation parameters, which are given in Supp. Table S20, and outputs a text file (.csv) of all pairwise Ks accumulated between all gene pairs (ohnologs) for each simulated genome. Architecturally, DemographiKS is designed as a pipeline with six modules, which are executed sequentially for the simulated polyploid. The module functions are (1) running SLiM, (2) initial paralog preparation, (3) gene shedding, (4) final paralog generation, (5) Ks calculation between paralogs, and (6) plotting the Ks histogram. We give details for each module in the “DemographiKS implementation” section of the supplemental methods.

### Running DemographiKS

DemographiKS is run by calling the “demographiKS.py” function from python and passing it the configuration xml file. For example, “python3 demographiKS.py myconfig.xml” DemographiKS currently includes no “hard” internal checks to verify that the ancestral population has achieved a steady state before continuing on to initiate parental divergence. This must be verified by the user, and details are given in the “The prerequisite of a steady-state” section of the supplementary methods.

### DemographiKS Output

The primary DemographiKS output file is a .csv file, with each line giving information for a unique gene-pair. The data are organized into four columns, giving the CODEML algorithm name used to calculate Ks, the names of the two genes (the genome ID and the paralog name), the Ks value, and the path to the CODEML output file from which the Ks value was extracted, respectively. This file is found in the “demographiKS_output” subfolder. By default, the file is named “[SimName].csv”. By default the simulation name is “allotetraploid_bottleneck” but this can be reset in the configuration file. The tree-sequence recording files with the genealogy data are output to the “SLiM_output” folder.

### Parallelization and replicates

Because DemographiKS is a forward-time genome- and population-level simulation, DemographiKS is computationally intensive. Due to random mating and recombination, DemographiKS is not easily parallelized. The work presented in this paper was done on a 16-node Dual-10 Xeon CPU (E5 −2630v4 2.20GHz) 15Tb RAM node (“goodmem”) on the mesxuuyan HPC at San Diego State University. Owing to the computationally intensive steps in simulating population-level and species-level processes, some simulations described here took more than 10 hours to complete. To facilitate running replicates, separate random seeds for SLim, MSPrime, and DemographiKS may be set in the configuration file. We recommend submitting each polyploid simulation as a separate batch job to a job scheduler such as SGE (Oracle) or SLURM (SchedMD).

### Using DemographiKS to model a genome with different homeologous exchange rates

To combine results from multiple DemographiKS runs, the output file containing the Ks values “allotetraploid_bottleneck.csv” from each run was concatenated.

### Simulating Ks data for paralogs maintained by selection

To simulate Ks data for paralogs maintained by selection, we derived a probability distribution based on the model set forth by Blanc and Wolfe 2004, and used python random.uniform() to sample the required number of Ks values (representing paralogous pairs) from the given distribution. The number of values generated was configurable and proportional to the number of Ks values output from DemographiKS. Our distribution for a random birth-death process with escape via selection is detained in the “Simulating Ks data for paralogs maintained by selection” section of the supplementary methods.

### Source code and data availability

DemographiKS is available from the Github repository https://github.com/tamsen/DemographiKS. DemographiKS v1.1.0.0 was used in this analysis, and input files used to drive the simulation and the scripts used to generate the figures are at https://github.com/tamsen/DemographiKS_paper_scripts.

### Transcriptomic Data

Empirical Ks histograms in Fig. 5 were generated from transcriptomic data for *Coffea arabica*, *Zea mays*, *Populus trichocarpa* and *Saccharum spontaneum.* Transcriptomic data were sourced from NCBI (NCBI assembly GCF_003713225.1, GCF_902167145.1, and GCA_022457205.1), and the software package KsRates (118) was used to calculate the Ks histograms for each species.

## Supporting information

Supplemental Methods and Discussion

## Acknowledgements

We would like to thank members of the Barker and Sethuraman Labs for valuable inputs during the conceptualization stage of this project. We would like to acknowledge Michael Barker, Matthew Hahn and Jonathan Wendel for valuable suggestions and spirited discussion. We would like to thank Ben Haller, Paul D Blischak, Mathews Sajan, Michael Barker, and Ryan Gutenkunst for their SLiM resources on modeling polyploids. All simulations were performed on the mesxuuyan HPC at San Diego State University, which was funded by NSF ABI: 1564659 to PI Sethuraman and coPI Jody Hey (Temple University) and startup funds to Sethuraman. TD and AS were both funded by NSF CAREER: 2147812 to PI Sethuraman, DE-SC0025673 to PI Archana Anand (SFSU) and coPI Sethuraman, and a CSUBIOTECH award to TD. This material is based upon work supported by the NSF Postdoctoral Research Fellowships in Biology Program under Grant No. 2507596. The funders had no role in this project’s development, design and delivery.

