## Supplemental Figures for "Hidden in Plain Sight. How Ks histogram dynamics can reveal and obscure ancient whole genome duplications"

**The DemographiKS software package**

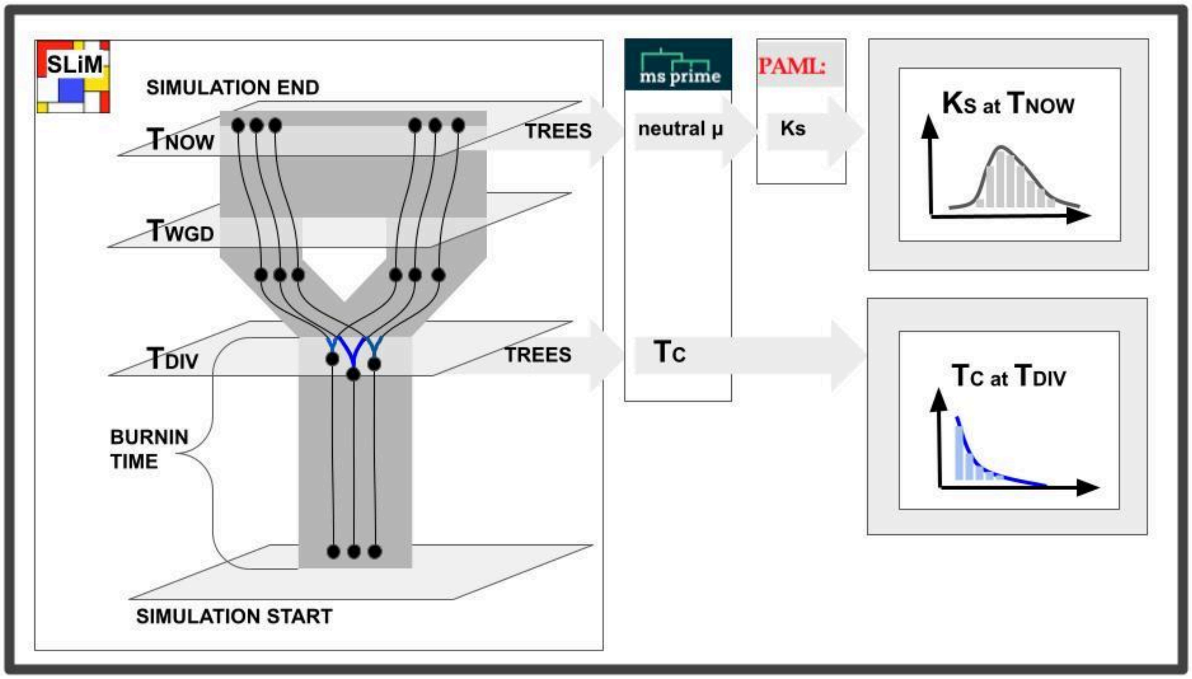


**Supp. Figure S1. The DemographiKS software package wraps three major components: SLiM, MSprime and PAML**. The polyploid simulation is based on SLiM scripts by (Blischak et al. 2023). This cartoon shows the SLiM internal model of allopolyploidy in the “SLiM” box on the left. The species tree is shown in gray, and the gene trees for three example genes are depicted by the black lines. The cartoon on the right shows how data exported from the SLiM model is used to derive the ancestral coalescent distribution (bottom right) and the polyploid Ks distribution (top right).The blue lines on the gene trees to the left highlight the time between the ancestral divergence (to become two diploid species) and coalescence (between two gene copies in the ancestral population), and the corresponding ancestral coalescent distribution is given in blue on the bottom right.

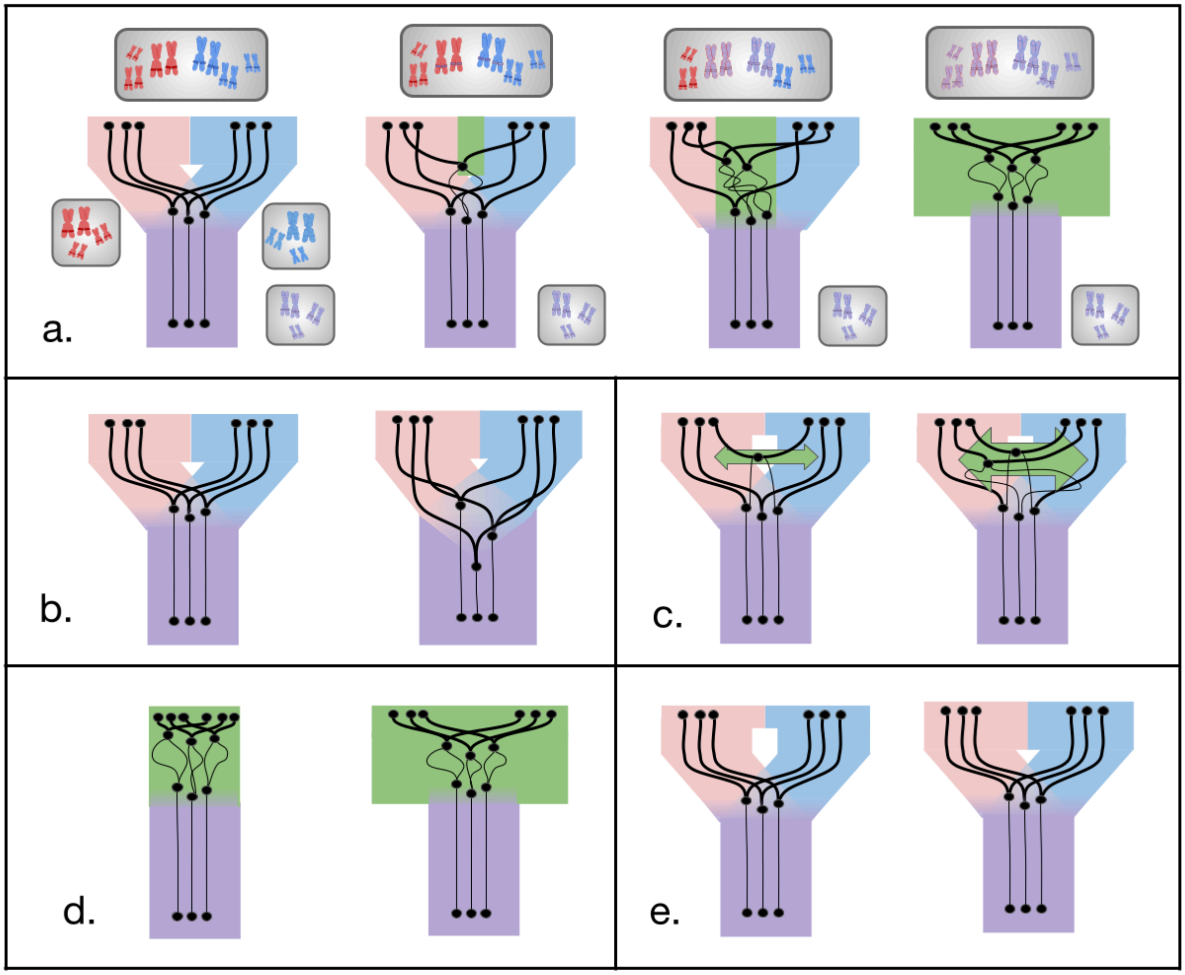


**Supp. Figure S2. A cartoon representation of the DemographiKS model.** The ancestral diploid species is shown in purple. The divergence of the ancestral diploid to form two separate species is illustrated by the separation of the pink and blue branches. Genetic exchange such as homeologous exchange and migration are shown in green. For an ancestral allopolyploid, the WGD event is illustrated by the pink and blue branches regaining contact with each other.
For an ancestral autopolyploid, the WGD event is illustrated by the purple diploid transitioning to a green color, signifying that exchange of genetic material is free and continuous between the increased multiplicity of homologous chromosome copies. The black lines indicate gene trees. The black circles represent nodes such as coalescent events on the gene trees.

**a. Increasing levels of homoeologous exchange.** On the far left of the first row, we give a cartoon representation of an ancestral allopolyploid with no genetic exchange between subgenomes (pink and blue). The timing of the Ks peak corresponds to the time of divergence of the ancestral diploids. The divergence of each individual paralogous pair corresponds to the divergence of the original orthologs at the time of divergence (black nodes) not the time of WGD. This model, with no homoeologous exchange, may be most appropriate for a young allopolyploid with highly differentiated subgenomes. Panels two and three demonstrate modeling for polyploids which have less differentiated subgnomes. Here we allow a moderate level of homeologous exchange (ie, a segmental polyploid whose closely-related parents would allow some small level of homeologous exchange) or a mix of homeologous exchange rates for different chromosomes (an allopolyploid with some gene conversion, or an autopolyploid shifting from tetrasomic to disomic inheritance). To model the pure autopolyploid (top row, last panel) we crank the homeologous exchange rate up all the way, to allow pure tetrasomic inheritance. DemographiKS still has labeled subgenomes (computationally speaking), but by that point for the autopolyploid, its biologically meaningless bookkeeping. With all the homeologous exchange, there are no subgenomes maintained. The formation of each gamete for each generation is N=4, choose 2. In all cases, to calculate the Ks distribution, we randomly take one chromosome per (labeled) subgenome per individual, and calculate the Ks between all ohnologous pairs. We note that for an autotetraploid, this is equivalent to randomly choosing any two chromosomes from the four available, since no subgenomes have been maintained in any way except by labeling.

**b. Increasing levels of ancestral diversity.** In the left box on the second row, we show how increasing levels of ancestral genetic diversity cause a greater diversity of coalescent times between ancestral orthologs. In the left panel in this box, we show an allopolyploid derived from an ancestral species with low levels of diversity, yielding a tighter coalescence. In the right panel of this box, we show an allopolyploid derived from an ancestral species with high levels of diversity. This yields a broader range of coalescent times, translating into a longer tail in the peak of the Ks histogram of the resulting allopolyploid.

**c. Increasing levels of migration.** In the right box of the second row, we show how increasing levels of migration between the parental diploids may cause some gene trees to have a coalescent time that is more recent than the time of parental divergence. In the left panel, we show how a small amount of gene flow between parental genomes may yield a small secondary peak in the Ks histogram of the resulting allopolyploid, closer to Ks=0 (left panel of this box). But as gene flow increases, the secondary Ks peak( due to migration) may eclipse the primary Ks peak (due to parental divergence) (right panel of this box).

**d. Increasing levels of diversity in the polyploid population.** In the left box of the third row, we show how increasing levels of genetic diversity in the autopolyploid yields greater diversity of coalescent times between homologs. In the left panel, we show that Ks will be identically zero for an autopolyploid with no diversity between its homologs. This model represents an autopolyploid that just came into being, from a highly inbred population, where the four homologous chromosome copies are identical. In the right panel, we consider an autopolyploid that results from the fusion of two individuals in the population, with slightly differentiated genomes. Thus we no longer have Ks=0 for all pairs. Instead the Ks distribution will have a peak at zero but mean at 4Ne (assuming panmixia and tetrasomic inheritance patterns).

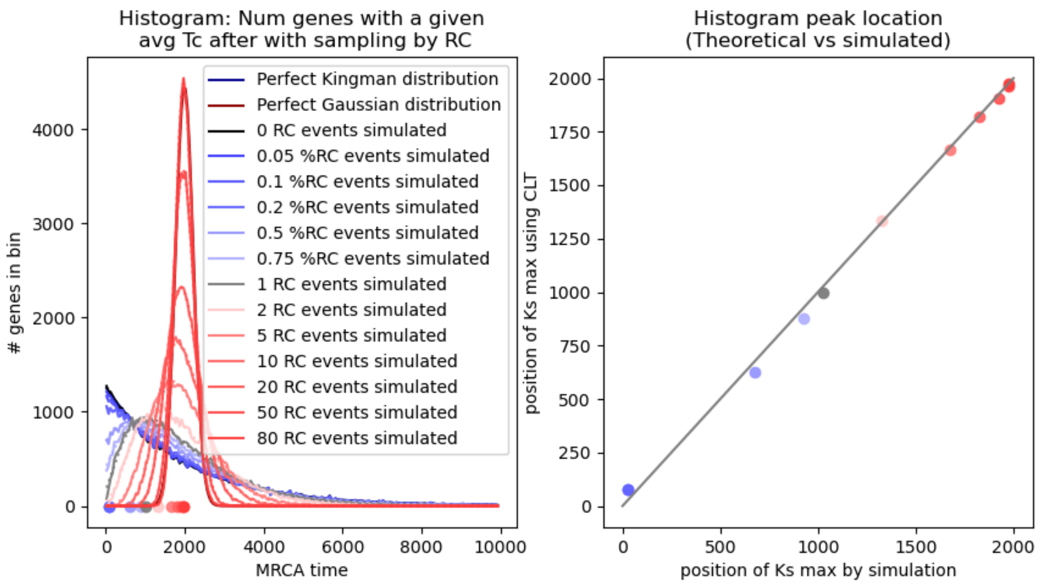


**Supp. Figure S3. Using a custom python script to directly resample the Kingman distribution, we show that the action of the central limit theorem is sufficient to recreate the diversity of Ks distributions shapes observed in the DemographiKS output.** Left: Distribution of coalescent times for 1000 genes in a test genome. Blue lines are a result of an increasing proportion of genes being the mixture of two different ancestral individuals' genomes. Red lines are the results of all genes being the mixture of an increasing number of different ancestral individuals' genomes. Right: The observed Ks peak by simulation and the predicted peak, based on the central limit theorem.

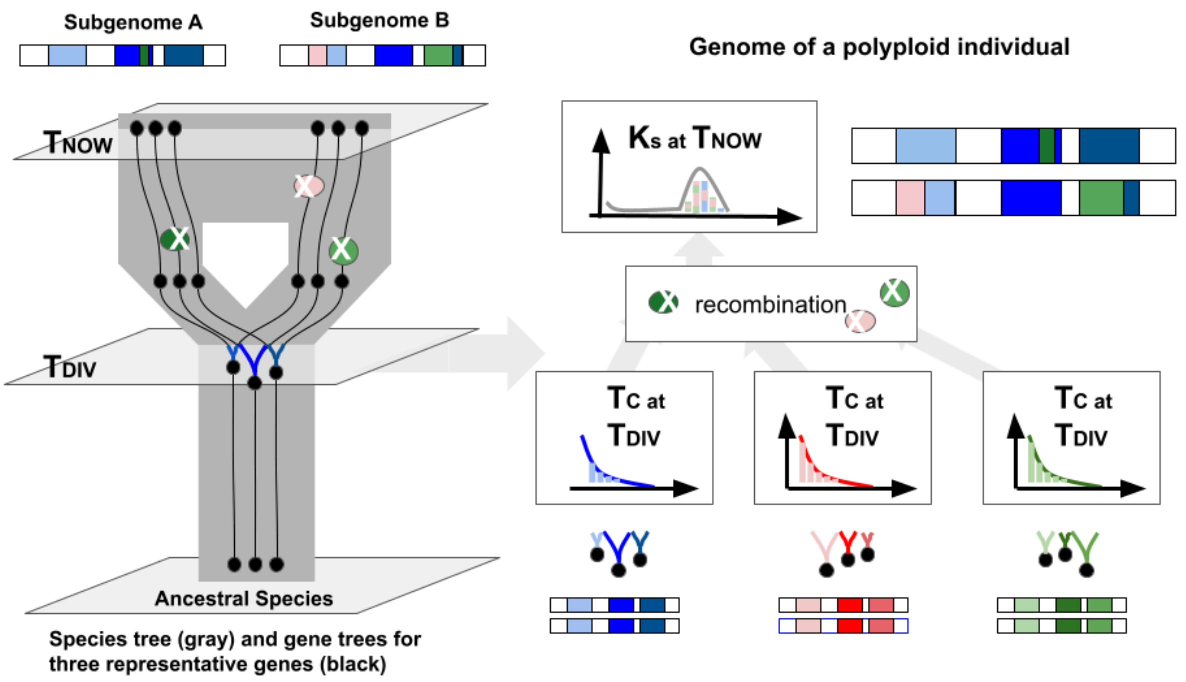


**Supp. Figure S4. A conceptual figure showing the Ks distribution for a single polyploid individual with recombination.** The species tree on the left places it in the context of polyploid evolution (gray), and for three example genes from the blue individual (black lines). The cartoons on the right show recombination resamples the ancestral Kingman coalescent distribution from three individuals, leading to a polyploid individual with a Gaussian Ks distribution.

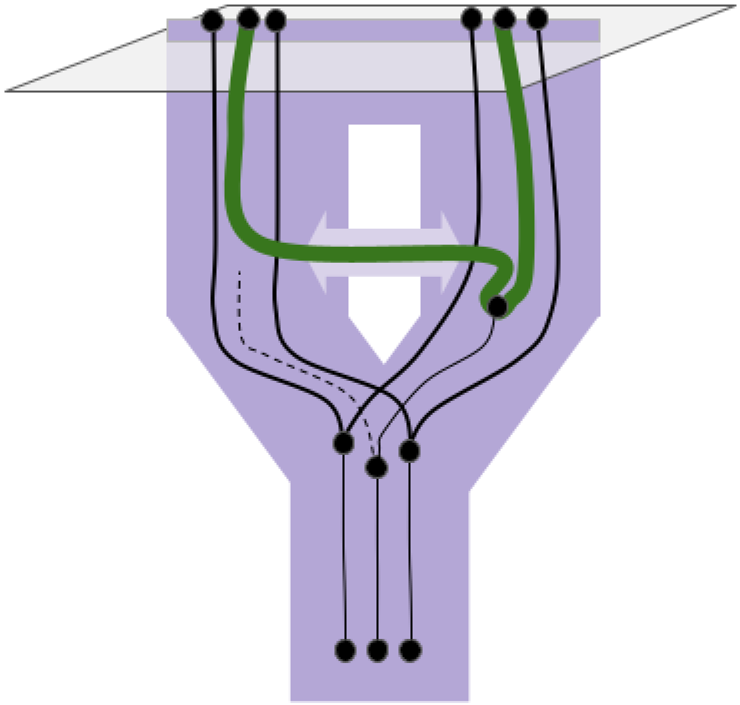

**Supp. Figure S5. A cartoon showing the Ks distribution given migration between the parental species.** Here the black subtrees represent gene trees within the purple species tree. The bifurcation represents ancestral divergence, when the genes become orthologs. After some time has elapsed the genes are reunited by whole genome duplication, and the gene pairs become paralogs. In this diagram one ancestral gene copy has been supplanted by gene flow due to migration. Thus the coalescence for the green gene pair is traced back to the migration event, not the parental divergence time.

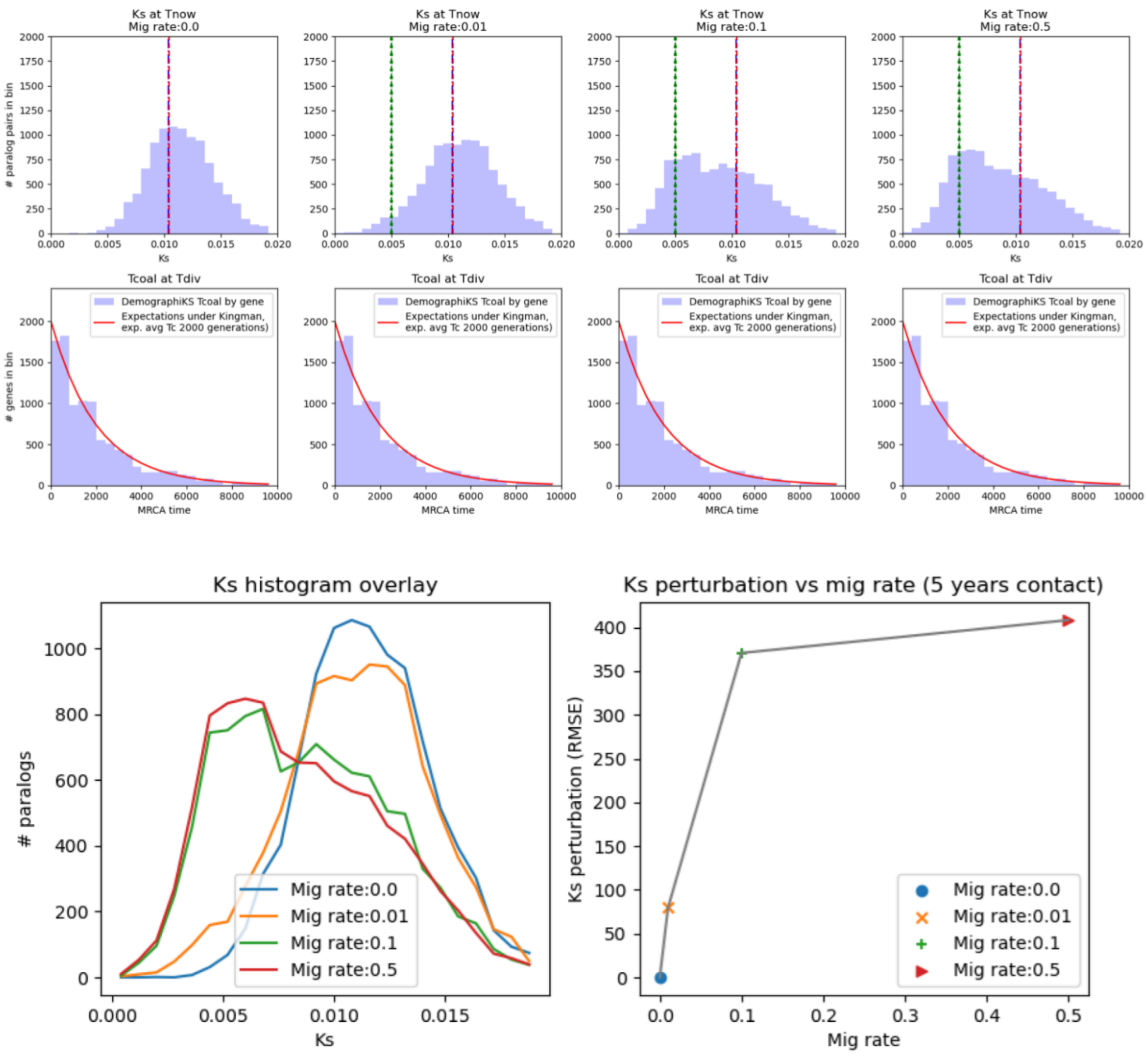


**Supp. Figure S6.** Five generations of secondary contact between the diploid parental species of the allopolyploid leaves a detectable signature in the Ks histogram, weak for low migration rates. Top: Ks histograms for migration rates from 0 to 50% over 5 generations. Bottom left, overlay of Ks histograms. The red dashed line indicates the predicted distribution center of mass, without migration. The green dashed line indicates the time of migration, in Ks-space. Bottom : Perturbations between the null hypothesis (no migration, blue) and the given migration rate Ks histogram (left), and measured as RMSE (right). Configuration parameters are given in Supp. Table S7.

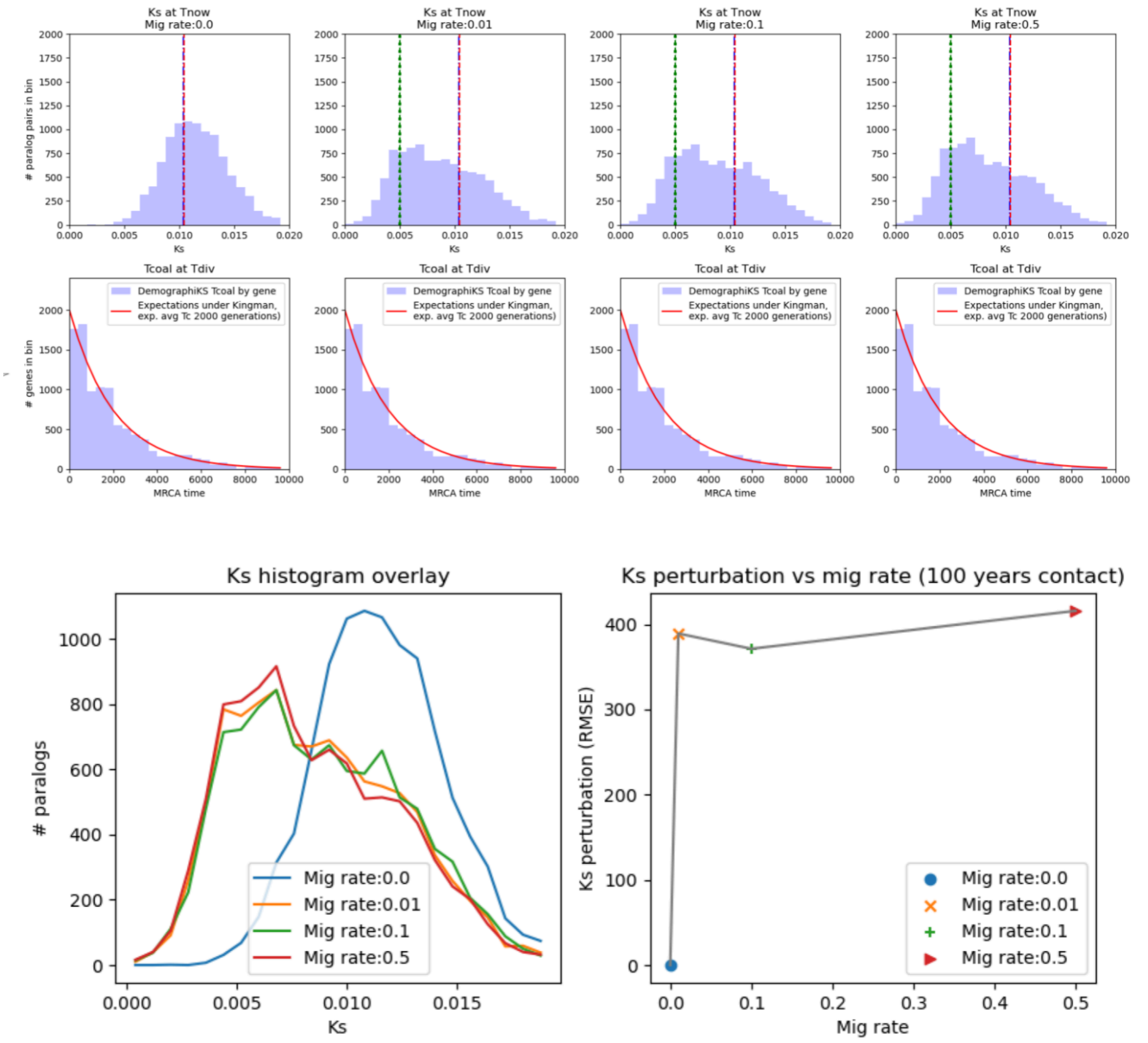


**Supp. Figure S7.** 100 generations of secondary contact between the diploid parental species of the allopolyploid leaves a strong signature in the Ks histogram, even for low migration rates. Top: Ks histograms for migration rates from 0 to 50% over 100 generations. Bottom left, overlay of Ks histograms. The red dashed line indicates the predicted distribution center of mass, without migration. The green dashed line indicates the time of migration, in Ks-space. Bottom : Perturbations between the null hypothesis (no migration, blue) and the given migration rate Ks histogram (left), and measured as RMSE (right). Configuration parameters are given in Supp. Table S7.

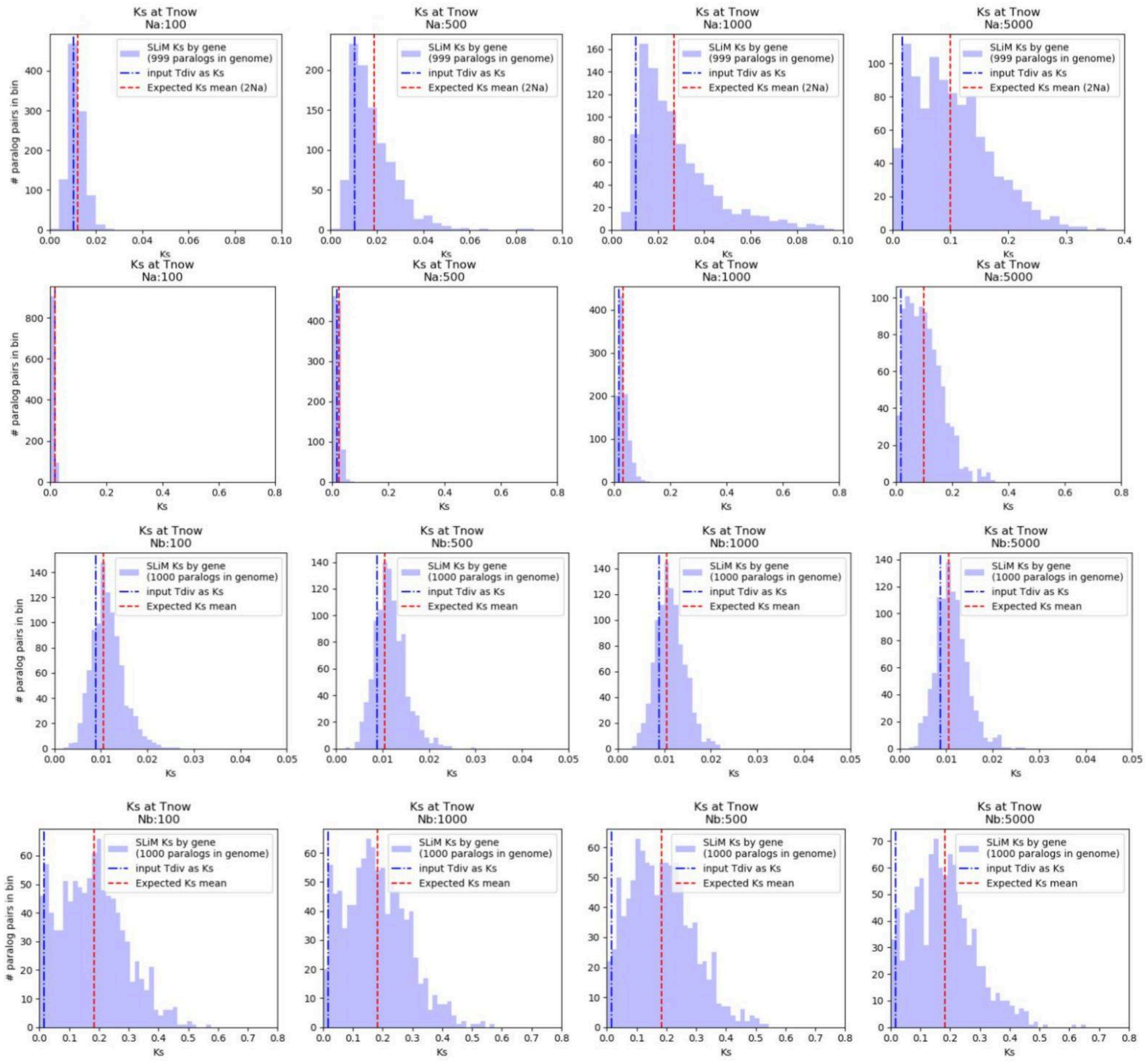


**Supp. Figure S8. The allopolyploid Ks histogram is influenced more strongly by ancestral population size than polyploid population size.** Top two rows: The allopolyploid Ks histogram shows a strong response to ancestral population size ($N_a$). In row 1, $N_b$ is held at a constant 100, and $N_a$ varies from 100 to 5000. In row 2, $N_b$ is held at a constant 10K and $N_a$ varies from 100 to 5000. Bottom two rows: The allopolyploid Ks histogram shows very little response to polyploid population size ($N_b$). In row 3, $N_a$ is held at a constant 100, and $N_b$ varies from 100 to 5000. In row 4, $N_a$ is held at a constant 10K and $N_b$ varies from 100 to 5000. Configuration parameters are given in Supp. Table S11.

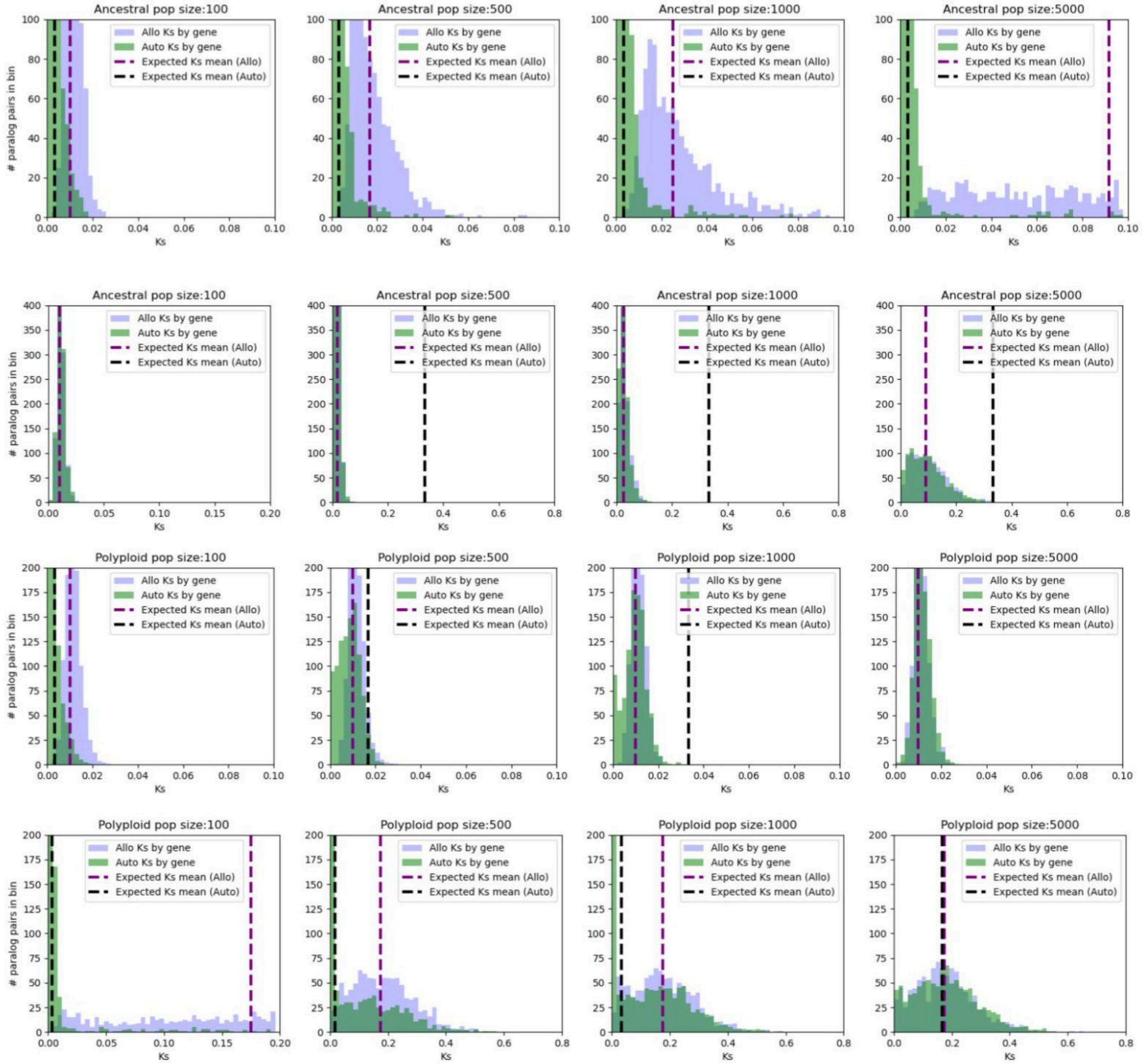


**Supp. Figure S9. The autopolyploid Ks histogram is influenced more strongly by polyploid population size than ancestral population size, unless polyploid population has recently expanded such that the theoretical polyploid coalescence (black dashed line) is further back in time than the coalescence of the ancestral diploid (purple dashed line), as shown in row 2.** Top two rows: Whether or not the autopolyploid Ks histogram responds to variation in ancestral population size ($N_a$) depends on polyploid population size ($N_b$). In row 1, $N_b$ is held at a constant 100, and $N_a$ varies from 100 to 5000. In row 2, $N_b$ is held at a constant 10K and $N_a$ varies from 100 to 5000. Bottom two rows: The autopolyploid Ks histogram always shows a strong response to polyploid population size ($N_b$). In row 3, $N_a$ is held at a constant 100, and $N_b$ varies from 100 to 5000. In row 4, $N_a$ is held at a constant 10K and $N_b$ varies from 100 to 5000. Configuration parameters are given in Supp. Table S11.

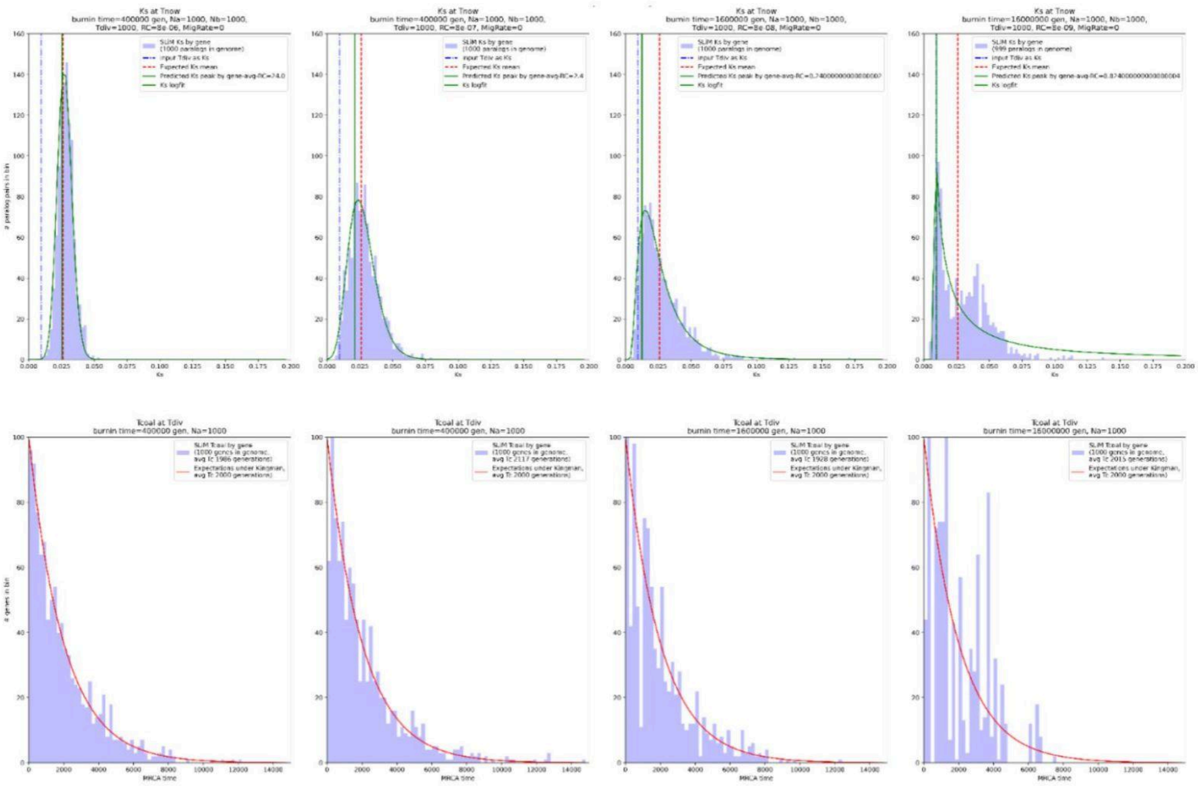

**Supp. Figure S10. A sparse ancestral coalescent (bottom row, right) can result in a lumpy Ks histograms (top row, left).** Top row gives the Ks histograms for four DemographiKS simulations of an allopolyploid. All simulations have the same fixed parameters, except for recombination rate, which varied from $8x10^{-6}$ to $8x10^{-9}$. Theory predicts that the Ks distributions should be a set of lognorms which appear more Gaussian on the left (due to more recombination) and more exponential on the right (due to less recombination). However, that trend fails in the right-most panel, where we see a lump due to artifacts caused by a sparse ancestral coalescent. Bottom row gives the distribution of ancestral coalescent times for the parental diploid, in generations, for the same four simulations described above. Theory predicts that the distributions will conform to an exponential (the Kingman coalescent). For distributions on the left, the high recombination rate is sufficient to break up linkage disequilibrium during the fixed burin period. For distributions on the right, especially in the last panel, the burnin is insufficient for the lower rates of recombination to break apart linkage: Thus many gene pairs have the same coalescent time, and the distribution is spotty and undersampled. As the simulation advances, this "lumpyness" propagates forward in time, causing artifacts in the final Ks distribution (top row, right).

### Effects of recombination rate on simulated Tc

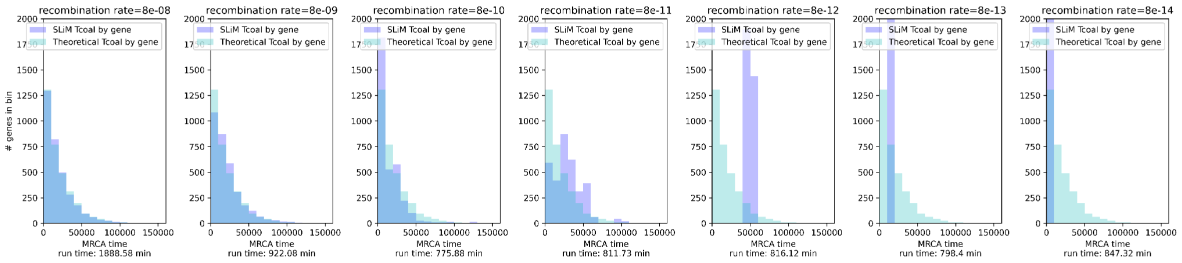


### Effects of Ne on simulated Tc

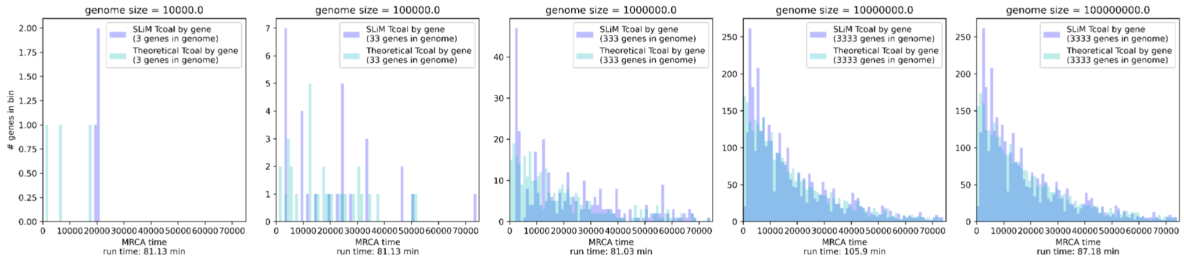


### Effects of genome size on simulated Tc Effects of burnin time on simulated Tc

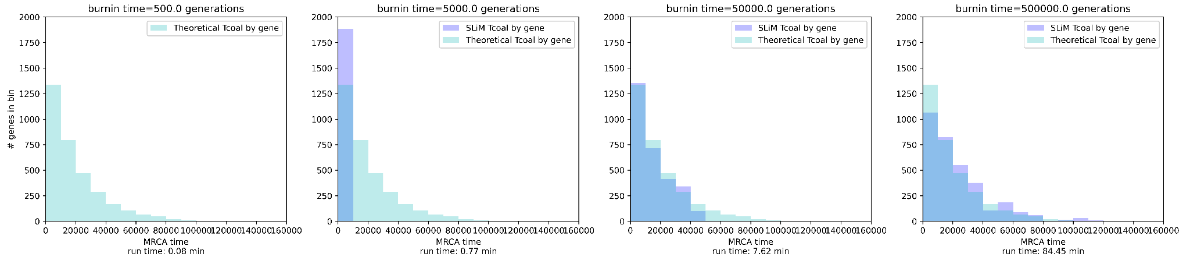


**Supp. Figure S11. Low genome size, low Ne and low recombination rates require more burn-in time to generate the expected Kingman distribution in the ancestral coalescent.** The plots above give the ancestral coalescent as determined by DemographiKS (purple), for a range of suboptimal parameters. Theoretical expectations based on subsampling alone are given in light blue.
