## Supplemental Methods and Discussion for "Hidden in Plain Sight. How Ks histogram dynamics can reveal and obscure ancient whole genome duplications"

#### DemographiKS implementation

DemographiKS is implemented as a pipeline application in python3. It wraps three major components: SLiM, MSprime, and PAML, as shown in Supp. Fig S1 (1–3). DemographiKS takes as input an XML configuration file listing the simulation parameters (Supp. Table S20) and outputs a text file (.csv) of all pairwise Ks accumulated between all gene pairs (ohnologs) for each simulated genome. Architecturally, DemographiKS is designed as a pipeline with six modules, which are executed sequentially for the simulated polyploid. The module functions are (1) running SLiM, (2) initial paralog preparation, (3) gene shedding, (4) final paralog generation, (5) Ks calculation between paralogs, and (6) plotting the Ks histogram. We give details for each module below.

**Step 1, running SLiM.** DemographiKS translates the input configuration file into arguments that are passed to SLiM (version 4.2.2). SLiM runs as an internal process which uses tree-sequence recording to save and export the population's genealogy (2). The SLiM scripts wrapped by DemographiKS are mechanistically identical to the scripts from (4), with the addition of Tree-sequence recording.

**Step2, initial paralog preparation.** DemographiKS uses tskit (version 0.5.6) to load the tree-sequence file generated by SLiM (5, 6). Within tskit, msprime is used to apply neutral mutations to the genealogy. We use the msprime default evolutionary model (msprime.JC69), which is a symmetrical mutation model. Once the final genomes have been appropriately populated with mutations, a random pair of focal genomes (one from each population of genomes representing the subgenomes) are selected to represent the genomes of the sampled polyploid, and the resulting paralogs are enumerated.

**Step 3, gene shedding.** Genes duplicated by WGD have a mean life expectancy given by an exponential decay distribution (7). The proportion of WGD duplicates which will be "dead" before the simulation concludes are calculated based on WGD time and pruned for computational efficiency. For traceability, pruned paralogs are recorded in the log file. Ohnolog life expectancy is configurable and based on (8, 9).

**Step 4, final paralog generation.** Genes which are not shed are written to disk as fasta files, in preparation for use as input by CODEML in step 5. To save space, these files are immediately erased after they are used, but this is configurable.

**Step 5, Ks calculation between paralogs.** Ks is calculated between all paralogous genes , using the CODEML program within PAML v4.10.7. At the end of this Ks calculation step, DemographiKS outputs a text file (.csv) of the pairwise Ks accumulated between all paralogs. Codon frequencies, transition/transversion rate and dN/dS were set to match those used in EVOLVER (1, 10) and recent Ks simulators (11, 12). The full CODEML configuration file is available to the user and included in the Github repository.

**Step 6, plotting the Ks histogram.** DemographiKS thereon generates a default Ks histogram output as a .png file. The histogram is not configurable and will always have exactly 50 bins. The provision of this histogram is only meant to give a confirmation of the run success. It is expected that the end users will use their own visualization tools to generate more elegant histograms.

**Simulating Ks data with some paralogs maintained by selection**

To simulate Ks data with some paralogs maintained by selection, we derived a probability distribution based on the model set forth by Blanc and Wolfe 2004, and used python3 random.uniform() to sample the required number of Ks values (representing paralogous pairs) from the given distribution. The number of values generated was configurable and proportional to the number of Ks values output from DemographiKS. Our distribution for a random birth-death process with escape via selection is detailed below:

Under a random birth-death process, the probability of a paralogous gene born, not yet being shed, forms an exponential decay distribution, with the mean life expectancy being the inverse of the decay rate.

$$P_{not\ yet\ shed}(t) = e^{-kt}$$

Thus, the chance of a paralogous gene being shed is

$$P_{shed}(t) = 1 - e^{-kt}$$

If some portion **α** of genes in the “fated to die” state are instead sufficiently beneficial to be maintained by selection, the chance of escaping death would be

$$P_{escape} = \boldsymbol{\alpha} \times P_{shed}$$

$$P_{remaining} = P_{not\ yet\ shed} + \boldsymbol{\alpha} * P_{shed}$$

Thus, the probability distribution for SSDs remaining in the genome should be

$$P_{remaining} = e^{-kt} + \boldsymbol{\alpha} * (1 - e^{-kt})$$

With some algebra,

$$P_{remaining} = \boldsymbol{\alpha} + e^{-kt}(1 - \boldsymbol{\alpha})$$

Which we can see gives the desired behaviour of an exponential decay near Ks=0, and a horizontal asymptote as Ks increases. Integrating from 0 to 4 in Ks-space gives the normalization constant necessary for a well-behaved probability distribution function.

$$C = 4\boldsymbol{\alpha} + \frac{\boldsymbol{\alpha}-1}{k}(e^{-4k} - 1).$$

In our simulations , we let the mean life expectancy of a SSD gene ($k$) be 3 million years and the rate of escaping death (**α**) be 0.2, based on a gene birth rate of 0.00162 per gene per million years (11), and a rate of duplicate retention ranging to 0.00035 to 0.0007 per gene, per million years (13).

It is unclear what the true value of **α** should be, since the true proportion of retained duplicates is a function of the distribution of fitness effects, the efficacy of selection, and several other parameters which might be lineage specific or vary over time. In practice, we found that the value of $\alpha$ was less significant than the overall number of genes randomly drawn from the distribution, which was set to best fit the observed asymptote (8, 13–17).

#### The prerequisite of a steady-state

For every result presented in this paper, we first verified that the ancestral population had achieved a steady state before parental divergence was initiated in the simulations. We ensured that the steady state was achieved by checking that the expected number of mutations had asymptotically reached theoretical expectations under a neutral model (an example script "diploid_snm.slim" is available on our Github site). We also ensured that linkage equilibrium had been achieved and that the coalescent time ($T_C$) matched theoretical expectations under neutrality. This was vital to our results, because deviations from equilibrium conditions - particularly a sparse or spotty coalescent distribution - would inject a "lumpiness" into the Ks histogram and distort results, potentially causing false peaks (Supp. Fig. S9). We also note that linkage equilibrium takes much longer to achieve than simply ensuring that all members of the population achieve coalescence within the burn-in window. For our purposes, we found that burn-in requirements were much longer than suggested by the SLiM manual and were highly dependent on recombination rate, genome size, and population size, and we give examples of this in Supp. Fig. S10.

DemographiKS currently includes no "hard" internal checks to verify that the ancestral population has achieved a steady state before continuing on to initiate parental divergence. However, the SLiM tree-sequence file specific to the ancestral population, and the related simulated coalescent distribution and theoretical coalescent distribution under the Kingman assumption are generated by DemographiKS before parental divergence is instantiated. These are automatically placed in the DemographiKS output folder for the user to examine for convergence. We note that there are use cases where a user might deliberately experiment with an ancestral population not in equilibrium, for example under a selective sweep, or to debug code without the computational overhead of a full burnin.

### Supplemental Discussion

#### Caveats

Despite the sensitivity of the Ks histogram to various evolutionary genomic and demographic parameters that we have demonstrated via extensive simulations using DemographiKS here, it is not a panacea. Much like how the power of a telescope rests on its ability to collect photons from distant sources, the power of the Ks histogram rests on its ability to collect mutations acquired long ago. A significant accumulation of synonymous mutations are needed to resolve the shape of distant effects on the ancient coalescent on current observed Ks histograms. If the ancestral coalescent is too shallow or the mutation rate too small, the number of mutations collected will be too small to be perceived. Specifically, the average number of mutations collected inside the ancestral coalescent is $2N_a$ multiplied by the mutation rate ($\mu$). If $T_{DIV}$ is too big, and $N_a$ and $\mu$ too small, these perturbations may be difficult to detect amidst the background mutation rate across time. It is possible that advanced methods may be developed which allow for the collection of mutations from more individuals or sister lineages, and this may serve to increase resolution.

We also acknowledge DemographiKS is slow compared to SpecKS. Population-level simulations are by nature, computationally intensive. In this paper the DemographiKS simulations in Part I and Part II were performed with settings chosen to elucidate a principle within a reasonable run time, not to model a specific polyploid lineage. To save computational time, small populations and short time scales were used wherever possible. However, the results may be mapped to larger populations and greater time scales with the use of Q-scaling, as described in (18–20). In the next generation of Ks simulators, we anticipate that Ks distributions could be simulated at the genome level, not the population level. This could be accomplished with the additional incorporation of novel theoretical models of polyploid evolution derived from population-level tools such as DemographiKS. These improvements would provide more rapid results for larger populations and longer time scales, without the need for Q-scaling.

An additional limitation of the DemographiKS model is that we presume that the rates of recombination, mutation and homoeologous exchange are constant over time and spatially along a chromosome. In reality, particularly within a polyploid genome, we expect that all these values will change over time, due to diploidization and general molecular evolutionary effects. For example, we might see homoeologous exchange rates for specific chromosomes within a genome go up, due to gene conversion, or down due to differentiation. For an ancestral autopolyploid, some chromosomes may shift from polysomic to disomic inheritance at different timescales, which may create quite complex Ks histograms (21, 22).

Additionally, in our empirical fits, we often utilized a lower recombination rate than was suggested in the literature, else the histograms would tend to Gaussians faster than expected. One explanation for this might be that our model of recombination did not take into account population structure, specifically isolation by distance, and non-random, assortative, or disassortative mating. Since in real populations, recombination would be happening between similar genomes, rather than between random genomes selected from within the entire population, we might expect that in simulation, recombination had a stronger effect than would be observed in nature. Additionally, the effects of the rates of various parameters changing over time will surely integrate over long timescales, such that many different evolutionary histories might create the same Ks histogram. Future work will iteratively need to incorporate this level of complexity into a new generation of polyploid simulation tools.
