## Supplemental Tables for "Hidden in Plain Sight. How Ks histogram dynamics can reveal and obscure ancient whole genome duplications"

Supplementary Tables

| **Supp. Table S1. DemographiKS parameters used for testing the effects of $N_e$ ie ($N_a$=$N_b$) Figure 1, row 1.** | | |
|---|---|---|
| **Parameter** | **Setting** | **Motivation** |
| Chromosome:<br>> total_num_bases<br>> num_codons_in_a_gene | Chromosome:<br>> 3000000<br>> 1000 | Set to the low end of reasonable values for plants to minimize runtime. Chr size ~ 1/10 Arabidopsis thaliana ch1, which is 30427671 pb. |
| > recombination_rate | > 1.26e-7 | Set high to achieve steady state within a reasonable burnin. ~100x A.thal rates. |
| Population:<br>> ancestral_ne<br>> bottleneck_ne | Population:<br>**> 10-1000**<br>**> 10-1000** | Set to the low end of reasonable values for plants to minimize runtime. Arabidopsis thaliana is at 10,000 $N_e$ |
| > burnin_time | > 400000 | As needed for to achieve steady state |
| SequenceEvolution:<br>> WGD_gene_half_life<br>> mutation_rate | Population:<br>> 31e-6<br>> 1.0e-5 | WGD half life set to parity with previous SpecKS simulations, mutation rate high to better show structure of exposed Tc |
| Speciation:<br>> DIV_time_Ge<br>> WGD_time_Ge | Speciation:<br>> 1000<br>> 1000 | This is more like effects after 0.1MY, since we increased rates of mut and RC x100. This choice shows dependence on $N_e$ quickly. |
| These sims saw run times from 1-15 min | | |

**Supp. Table S2. SpecKS parameters used for testing the effects of $N_e$, Figure 1, row 1.**

| **Parameter** | **Setting** | **Motivation** |
|---|---|---|
| GeneTree:<br>> num_gene_trees<br>> mean_gene_birth_rate<br>> SSD_half_life_MY<br>> WGD_half_life_MY> | Chromosome:<br>> 1000<br>> 0<br>> 1<br>> 31e-2 | Set to match DemographiKS. Since DemographiKS does not include a gene birth/death model, gene birth rate is set to zero. The SSD half life is irrelevant since no SSDs will be born. WGD set to parity. |
| > recombination_rate | **> 0** | Trivially. Specks has no RC model |
| Polyploid:<br>> gene_div_time_dist._param<br>> $T_{DIV}$ & $T_{WGD}$ | Polyploid:<br>> expon,0, 0.00002-0.02<br>> 0.00001 | The genetree distribution is set to match the Kingman, thus describes an exponential dist. with rate parameter k=0.02=2*Ne/10^6 (to convert to MYA) |
| SequenceEvolution:<br>> Ks_per_Myr<br>> per_site_ev'n_distance | SequenceEv'n:<br>> 0.58333333<br>> 0.73977282 | > mutation rate = 7.0e-7> syn mut rate of 0.58333333 (ratio of 1.2 mut/synon.mut)<br>> for Ks-per-year at 0.01, ev'n distance is 0.01268182 (Tiley et al. 2018). We scale up. |
| SpecKS runtime ~2 min | | |

**Supp. Table S3. DemographiKS parameters used for testing the effects of $T_{DIV}$, Figure 1, row 2.**

| **Parameter** | **Setting** | **Motivation** |
|---|---|---|
| Chromosome:<br>> total_num_bases<br>> num_codons_in_a_gene | Chromosome:<br>> 10000000<br>> 1000 | Set to the low end of reasonable values for plants to minimize runtime. Chr size ~ 1/3 Arabidopsis thaliana<br>(Yields 3333 genes. 1000 is sufficient.) |
| > recombination_rate | > 8e-9 | Compromise between *Arabidopsis thaliana* at 8e-10 and a reasonable burnin |
| Population:<br>> ancestral_ne<br>> bottleneck_ne | Population:<br>> 1000<br>> 1000 | Set to the low end of reasonable values for plants to minimize runtime. Similar to values given for Arabidopsis thaliana |
| > burnin_time | > 5e7 | As needed for to achieve steady state |
| SequenceEvolution:<br>> WGD_gene_half_life<br>> mutation_rate | Population:<br>> 31e-6<br>> 1.2e-8 | Set to parity with previous SpecKS simulations |
| Speciation:<br>> DIV_time_Ge<br>> WGD_time_Ge | Speciation:<br>**> 10K-1MYA**<br>**> 10K-1MYA** | Set to parity with previous SpecKS simulations. Note DemographiKS units are years/generations, not MY. |
| These sims saw run times from 712 - 583 min, ~ 11 hours a run | | |

| **Supp. Table S4. SpecKS parameters used for testing the effects of $T_{DIV}$, Figure 1, row 2.** | | |
|---|---|---|
| **Parameter** | **Setting** | **Motivation** |
| GeneTree:<br>> num_gene_trees<br>> mean_gene_birth_rate<br>> SSD_half_life_MY<br>> WGD_half_life_MY> | Chromosome:<br>> 1000<br>> 0<br>> 1<br>> 31e-2 | Set to match DemographiKS. Since DemographiKS does not include a gene birth/death model, gene birth rate is set to zero. The SSD half life is irrelevant since no SSDs will be born. WGD set to parity. |
| > recombination_rate | > 0 | Trivially. Specks has no RC model |
| Polyploid:<br>> gene_div_time_dist._param<br>> $T_{DIV}$ & $T_{WGD}$ | Polyploid:<br>> expon,0,0.02<br>**>0.01-1.0 MY** | The genetree distribution is set to match the Kingman, thus describes an exponential dist. with rate parameter k=0.02=2*$N_e$/10^6 (to convert to MYA) |
| SequenceEvolution:<br>> Ks_per_Myr<br>> per_site_ev'n_distance | SequenceEv'n:<br>> 0.58333333<br>> 0.73977282 | > mutation rate = 7.0e-7-> syn mut rate of 0.58333333 (ratio of 1.2 mut/synon.mut)<br>> for Ks-per-year at 0.01, ev'n distance is 0.01268182 (Tiley et al. 2018). We scale up. |
| SpecKS runtime 13.3- 14.3 min | | |

| **Supp. Table S5. DemographiKS parameters used for testing recombination, Figure 1, row 3.** | | |
|---|---|---|
| **Parameter** | **Setting** | **Motivation** |
| Chromosome:<br>> total_num_bases<br>> num_codons_in_a_gene | Chromosome:<br>> 3000000<br>> 1000 | Set to the low end of reasonable values for plants to minimize runtime. Chr size ~ 1/10 *Arabidopsis thaliana* |
| > recombination_rate | **> 8e-10 - 8e-6** | Set to exceed/span reasonable values, to test edge case behaviors. Includes 100x RC rate increase for plants to match all time-dependent parameters. |
| Population:<br>> ancestral_ne<br>> bottleneck_ne | Population:<br>> 10000<br>> 10000 | Set to the low end of reasonable values for plants to minimize runtime. SpecKS simulations covered $N_e$ up to $10^6$ |
| > burnin_time | > 4e3-4e6 | As needed for to achieve steady state |
| SequenceEvolution:<br>> WGD_gene_half_life<br>> mutation_rate | Population:<br>> 31e-4<br>> 7.0e-7 | ~100x common rates for plants, to speed up results generation. (A.thal. Has a mutation rate of $7e^{-9}$ ). RC also at 100x. |
| Speciation:<br>> DIV_time_Ge<br>> WGD_time_Ge | Speciation:<br>> 10<br>> 10 | Set low to minimize runtime.<br>Effects are measurable right at $T_{DIV}$.<br>(And most clear at that time) |
| These sims saw run times from 695,29,67,579 min, ~ max 11 hours | | |

| **Supp. Table S6. SpecKS parameters used for testing recombination, Figure 1, row 3.** | | |
|---|---|---|
| **Parameter** | **Setting** | **Motivation** |
| GeneTree:<br>> num_gene_trees<br>> mean_gene_birth_rate<br>> SSD_half_life_MY<br>> WGD_half_life_MY> | Chromosome:<br>> 1000<br>> 0<br>> 1<br>> 31e-2 | Set to match DemographiKS. Since DemographiKS does not include a gene birth/death model, gene birth rate is set to zero. The SSD half life is irrelevant since no SSDs will be born. WGD set to parity. |
| > recombination_rate | **> 0** | Trivially. Specks has no RC model |
| Polyploid:<br>> gene_div_time_dist._param<br>> $T_{DIV\&}T_{WGD}$ | Polyploid:<br>> expon,0,0.02<br>> 0.00001 | The genetree distribution is set to match the Kingman, thus describes an exponential dist. with rate parameter k=0.02=2*Ne/$10^6$ (to convert to MYA) |
| SequenceEvolution:<br>> Ks_per_Myr<br>> per_site_ev'n_distance | SequenceEv'n:<br>> 0.58333333<br>> 0.73977282 | > mutation rate = 7.0e-7-> syn mut rate of 0.58333333 (ratio of 1.2 mut/synon.mut)<br>> for Ks-per-year at 0.01, ev'n distance is 0.01268182 (Tiley et al. 2018). We scale up. |
| SpecKS runtime 2.98 min | | |

**Supp. Table S7. DemographiKS parameters used for testing the effects of migration, Supp. Fig. S5 (5 generations) and S6 (100 generations), and Figure 2 of main text (gradual speciation)**

| Parameter | Setting | Motivation |
|---|---|---|
| Chromosome:<br>> total_num_bases<br>> num_codons_in_a_gene | Chromosome:<br>> 30000000<br>> 1000 | Same as Arabidopsis thaliana chr 1 |
| > recombination_rate | > 8e-9 | Includes 10x RC rate increase for plants to match all time-dependent parameters.<br>Achieves equilibrium faster |
| Population:<br>> ancestral_ne (Na)<br>> bottleneck_ne (Nb) | Population:<br>> 1000<br>> 1000 | Na set such that Tc from Na << than Tc from Nb, so that the effects of Nb can be isolated from effects of Na |
| > burnin_time | > 10^8 | As needed for to achieve steady state |
| SequenceEvolution:<br>> WGD_gene_half_life<br>> mutation_rate | Population:<br>> 31e-6<br>> 1.2e-8 | Parity with SpecKS settings |
| Speciation:<br>> DIV_time_Ge<br>> WGD_time_Ge | Speciation:<br>> 10^6<br>> 1000 | WGD time set to be recent. This experiment isn't about gene shedding, and we don't want to shed genes we took so much time to simulate. |
| Migration (5yrs):<br>> mig_start_gen<br>> mig_stop_gen<br>> mig_rate | Migration:<br>> 500000<br>> 500005<br>> [0,0.01,0.1,0.5] | Test the effects of 5 years of migration at varying levels. Set out 500K from parental div, so the peak due to migration couldn't would stand out from the peak due to $T_{DIV}$ |
| Migration (100yrs):<br>> mig_start_gen<br>> mig_stop_gen<br>> mig_rate | Migration:<br>> 500000<br>> 500100<br>> [0,0.01,0.1,0.5] | Test the effects of 100 years of migration at varying levels. Set out 500K from parental div, so the peak due to migration couldn't would stand out from the peak due to $T_{DIV}$ |
| Migration (gradual):<br>> mig_start_gen<br>> mig_stop_gen<br>> mig_rate | Migration:<br>> 1<br>> [50K,100K, 250K,500K]+1<br>> [0,0.01,0.1,0.5] | Test the effects of continual migration directly following divergence, for varying rates and durations |
| Run time ~1260 min, 21 hours | | |

| **Supp. Table S8. DemographiKS parameters used for testing the effects of homoeologous exchange given varying population sizes, Figure 3, top row.** | | |
|---|---|---|
| **Parameter** | **Setting** | **Motivation** |
| Chromosome:<br>> total_num_bases<br>> num_codons_in_a_gene | Chromosome:<br>> 3000000<br>> 1000 | Set to the low end of reasonable values for plants to minimize runtime. Chr size ~ 1/10 *Arabidopsis thaliana* |
| > recombination_rate | > 8e-8 | Includes 100x RC rate increase for plants to match all time-dependent parameters. Achieves equilibrium faster, so can get by with less burnin time (sim still takes 1hr+) |
| Population:<br>> ancestral_ne (Na)<br>> bottleneck_ne (Nb) | Population:<br>> 10000<br>**> [50 -1000]** | $N_a$ set such that Tc from $N_a$ << than $T_c$ from $N_b$, so that the effects of $N_b$ can be isolated from effects of $N_a$ |
| > burnin_time | > 4e5 | As needed for to achieve steady state |
| SequenceEvolution:<br>> WGD_gene_half_life<br>> mutation_rate | Population:<br>> 31e-6<br>> 7.0e-5 | Mut rate set high and gene shedding set low for computational efficiency. (We want to clearly see the effects of $N_b$). |
| Speciation:<br>> DIV_time_Ge<br>> WGD_time_Ge | Speciation:<br>> False<br>> 5000 | No parental divergence since this is an autopolyploid. WGD time set high so that if the Ks histogram had a time dependency, it would be obvious. |
| Run time ~2 hours each | | |

**Supp. Table S9. DemographiKS parameters used for testing the effects of homoeologous exchange given varying time since WGD, Figure 3, bottom row.**

| **Parameter** | **Setting** | **Motivation** |
|---|---|---|
| Chromosome:<br>> total_num_bases<br>> num_codons_in_a_gene | Chromosome:<br>> 10000000<br>> 1000 | Set to the low end of reasonable values for plants to minimize runtime. Chr size ~ 1/3 *Arabidopsis thaliana*<br>(gave 3333 genes, probably more than I needed) |
| > recombination_rate | > 8e-9 | Compromise between *Arabidopsis thaliana* at 8e-10 and a reasonable burnin |
| Population:<br>> ancestral_ne<br>> bottleneck_ne | Population:<br>> 1000<br>> 1000 | Set to the low end of reasonable values for plants to minimize runtime. Similar to values given for *Arabidopsis thaliana* |
| > burnin_time | > 5e7 | As needed for to achieve steady state |
| SequenceEvolution:<br>> WGD_gene_half_life<br>> mutation_rate | Population:<br>> 31e-6<br>> 1.2e-8 | Set to parity with previous SpecKS simulations |
| Speciation:<br>> DIV_time_Ge<br>> WGD_time_Ge | Speciation:<br>**> 10K-1MYA (or False)**<br>**> 10K-1MYA** | Set to parity with previous SpecKS simulations. DIV_time_Ge set to “False” for autopolyploid runs. This turns off homoeologous recombination. |

Sims saw run times from 712 - 583 min, ~ 11 hours a run

**Supp. Table S10. Expectations and observations of changes/lack of changes in the simulated Ks histogram due to variation in Na and Na.**

| **Expections** | Test 1 | Test 2 | Test 3 | Test 4 |
|---|---|---|---|---|
| | Na varies from 100 to 5000 | | Nb varies from 100 to 5000 | |
| | Nb=100 | Nb=10K | Na=100 | Na=10K |
| No Hom Ex | ++ | ++ | -- | -- |
| Free Hom Ex | -- | ?? | ++ | ++ |

| **Observations** | Test 1 | Test 2 | Test 3 | Test 4 |
|---|---|---|---|---|
| | Na varies from 100 to 5000 | | Nb varies from 100 to 5000 | |
| | Nb=100 | Nb=10K | Na=100 | Na=10K |
| No Hom Ex | ++ | ++ | -- | -- |
| Free Hom Ex | -- | ++ | ++ | ++ |

**Top: We expected that allopolyploid Ks histogram shapes would be more affected by variation in ancestral population size ($N_a$) and that autopolyploid Ks histogram shapes would be more affected by variation in polyploid population size ($N_b$).**
**Bottom: Our results showed unexpected coupling between the effects of $N_a$ and $N_b$ for autopolyploids for higher $N_b$.** Here "++" or "--" denotes whether the simulated Ks histogram shows/expected to show strong ("++") or weak ("--") dependency on the test variable ($N_a$ or $N_b$). "??" denotes anticipated coupling between $N_a$ and $N_b$.

| **Supp. Table S11. DemographiKS parameters used to test the effects of variations in $N_a$ and $N_b$, when homeologous exchange is (a) not allowed and (b) allowed. Figure 7.** | | |
|---|---|---|
| **Parameter** | **Setting** | **Motivation** |
| Chromosome:<br>> total_num_bases<br>> num_codons_in_a_gene | Chromosome:<br>> 3000000<br>> 1000 | Set to the low end of reasonable values for plants to minimize runtime. Chr size ~ 1/3 *Arabidopsis thaliana*<br>(gave 3333 genes, probably more than I needed) |
| > recombination_rate | > 8e-8 | Compromise between *Arabidopsis thalian*a at 8e-10 and a reasonable burnin |
| > homoeologous_exchange_ rate | (a) 0.0<br>(b) 0.5 | Testing the effects of homoeologous exchange. |
| Population:<br>> ancestral_ne<br>> bottleneck_ne | Population**:**<br>**> 100 or 10K**<br>**> [100 - 5000]** | Set to span the low end of reasonable values, to elucidate the dependence on the polyploid populations size $N_b$ |
| > burnin_time | > 5e5 | As needed for to achieve steady state |
| SequenceEvolution:<br>> WGD_gene_half_life<br>> mutation_rate | Population:<br>> 31e-6<br>> 1.0e-5 | WGD_gene_half_life set to parity with previous SpecKS simulations. Mutation rate fast to elucidate changes in $N_b$ |
| Speciation:<br>> DIV_time_Ge<br>> WGD_time_Ge | Speciation:<br>> 1000<br>> 1000 | Set to parity with previous SpecKS simulations. |
| Sims saw run times from 5 min - 1 hour | | |

**Supp. Table S12. Parameter estimates for *Coffea arabica* from the literature**

| | | | |
|---|---|---|---|
| chromosome | total_num_bases | 2.88 M in CDS for original Chr 7 | https://www.ncbi.nlm.nih.gov/gdv/browser/genome/?id=GCF_003713225.1 , https://www.nature.com/articles/s41467-023-44449-8 |
| | num_codons_in_a_gene | 800 | "A chromosome-scale assembly reveals chromosomal aberrations and exchanges generating genetic diversity in *Coffea arabica* germplasm |
| | recombination_rate | 1 × 10−8..? | 1 × 10−8 is used in "A single polyploidization event at the origin of the tetraploid genome of Coffea arabica is responsible for the extremely low genetic variation in wild and cultivated germplasm"<br>(Arabidopsis, 8.06452e-10; Rice, 8.97e-10, Helianthus, 4e-09) |
| | homeologous exchange rate | 0 | Genetic Diversity of Arabica Coffee (*Coffea arabica L.*) in Nicaragua as Estimated by Simple Sequence Repeat Markers<br>""The results clearly suggest the lack of recombination between the chromosomes of the two ancestral genomes due to the amphidiploid nature of arabica coffee." |
| Population | Na | 10^5-10^6 | "The genome and population genomics of allopolyploid Coffea arabica reveal the diversification history of modern coffee cultivars" |
| | Nb | 10^4 | "between 10,000 and 50,000 individuals"<br><br>The genome and population genomics of allopolyploid *Coffea arabica* reveal the diversification history of modern coffee cultivars<br><br>Newly sequenced genome reveals coffee's prehistoric origin story, and its future under climate change |
| Seq. Evolution | WGD_gene_half_life | 3e7 | Accurate Inference of the Polyploid Continuum Using Forward-Time Simulations 2024 |
| | mutation_rate | 7.77 x 10-9 | "The genome and population genomics of allopolyploid *Coffea arabica* reveal the diversification history of modern coffee cultivars" ". Using a mutation rate of 7.77 x 10-9 / (bp*generation)" |

| | | | |
|---|---|---|---|
| | | | https://www.biorxiv.org/content/10.1101/2023.09.06.556570v1.full |
| Speciation | DIV time | 1.44 - 0.675 MYA | The genome and population genomics of allopolyploid Coffea arabica reveal the diversification history of modern coffee cultivars<br><br>“We find evidence for a founding polyploidy event 350,000–610,000 years ago, followed by several pre-domestication bottlenecks |
| | WGD time | 0.35 - 0.610 MYA | |

**Supp. Table S13. Parameter estimates for *Zea mays* from the literature**

| | | | |
|---|---|---|---|
| chromosome | total_num_bases | 152M bp, origins CDS 11.4M | ch 10 is the shortest. NCBI GCF_902167145.1, Sequence composition and genome organization of maize” |
| | num_codons_in_a_gene | 1500 | Uneven chromosome contraction and expansion in the maize genome. Genome Res. 2006 |
| | recombination_rate | 7.3*10^-9 | “Intraspecific variation of recombination rate in maize” 2013 |
| | homeologous exchange rate | Segmental. Varies across chr, 0-0.5 | “Maize as a model for the evolution of plant nuclear genome”<br>“Study on Haploid Inducing and Its Meiotic Abnormality in Maize” |
| Population | Na | 10^5 | “The interplay of demography and selection during maize domestication and expansion” |
| | Nb | 1000 (bottleneck) max at 10^7 | “The interplay of demography and selection during maize domestication and expansion”<br>“Fast diffusion of domesticated maize to temperate zones” |
| Seq. Evolution | WGD_gene_half_life | 5-31 My | Gene Loss and Movement in the Maize Genome 2004 + Accurate Inference of the Polyploid Continuum Using Forward-Time Simulations 2024 |
| | mutation_rate | 2.17-3∗10−8 | 3.0 The interplay of demography and selection during maize domestication and expansion<br>2.17 “Contributions of *Zea mays* subspecies mexicana haplotypes to modern maize” |
| Speciation | DIV time | 20.5 mya<br>11.9 Mya | “Maize as a model for the evolution of plant nuclear genomes - Gaut 2000<br>Physical and genetic structure of the maize genome reflects its complex evolutionary history.<br>Maize and wild relatives show distinct patterns of genome downsizing following polyploidy (5-12) |
| | WGD time | 11.4 Mya<br>5-12 Mya | |

**Supp. Table S14. Parameter estimates for *Populus trichocarpa* from the literature**

| | | | |
|---|---|---|---|
| chromosome | total_num_bases | 13,000,000 | NCBI. Chr 6 of 19 https://www.ncbi.nlm.nih.gov/gdv/browser/genome/?id=GCF_000002775.5 |
| | num_codons_in_a_gene | 1300 | ~4000 bp per gene = 1300 codons per gene https://www.ncbi.nlm.nih.gov/refseq/annotation_euk/Populus_trichocarpa/101/ |
| | recombination_rate | (4.2-7.8 )*10^-8 | “High-resolution mapping reveals hotspots and sex-biased recombination in Populus trichocarpa” |
| | homeologous exchange rate | Probably varies for diff chr, but low by now | Expect chr are pretty well-differentiated after ~58 MY since salicoid WGD event |
| Population | Na | ??? | Unclear. Probably Na > Nb. We don’t know. |
| | Nb | 4000–6000 | Genome resequencing reveals multiscale geographic structure and extensive linkage disequilibrium in the forest tree Populus trichocarpa |
| Seq. Evolution | WGD_gene_half_life | 31 MY | Accurate Inference of the Polyploid Continuum Using Forward-Time Simulations 2024 |
| | mutation_rate | (4.2–5.2)×10^-8 | A genome assembly and the somatic genetic and epigenetic mutation rate in a wild long-lived perennial Populus trichocarpa |
| Speciation | DIV time | ~4*10^6 | The Salicoid WGD occurred 58 MYA, ~4*10^6 generations. “The Ancient Salicoid Genome Duplication Event: A Platform for Reconstruction of De Novo Gene Evolution in Populus trichocarpa” |
| | WGD time | ~4*10^6 | |

| **Supp. Table S15. Parameter estimates for *Saccharum spontaneum* from the literature** | | | |
|---|---|---|---|
| chromosome | total_num_bases | chr varies from 100-44 MB | NCBI https://www.ncbi.nlm.nih.gov/datasets/genome/?taxon=62335 |
| | num_codons_in_a_gene | 1000 | Simplifying assumption from Tiley 2018 Assessing the Performance of Ks “Plots for Detecting Ancient Whole Genome Duplications” |
| | recombination_rate | 8.97e-10 (Rice) ~1e-9 | StdPopSim catalog, borrowed recombination rate for Rice |
| | homeologous exchange | Frequent | The complex polyploid genome architecture of sugarcane |
| Population | Na | > e^6? | Predicting invasion risk of grasses in novel environments requires improved genomic understanding of adaptive potential |
| | Nb | 0.01-0.5e6 | Genomic insights into the recent chromosome reduction of autopolyploid sugarcane Saccharum spontaneum |
| Seq. Evolution | WGD_gene_half_life | 30 *10^6 | Simplifying assumption from Dunn Accurate Inference of the Polyploid Continuum Using Forward-Time Simulations |
| | mutation_rate | $3.8 \times 10{-}9$ | The complex polyploid genome architecture of sugarcane |
| Speciation | DIV time | 1.5 MY, sugar WGD, 70 MY ancient cereals WGD | Genomic insights into the recent chromosome reduction of autopolyploid sugarcane Saccharum spontaneum |
| | WGD time | | |

**Supp. Table S16. DemographiKS parameters for *Coffea arabica***
All genes w/o HomEx, run name EMP_Coff_42_m09d19y2025_h11m47s34

| | | Est. from prior works | DGKS Q1 (theory) | DGKS Q100 (run 42) |
|---|---|---|---|---|
| chromosome | total_num_bases | 2.88 M in CDS | 3000000 | 3000000 |
| | num_codons_in_a_gene | 800 | 800 | 800 |
| | recombination_rate | 1 × 10−8..? (1 × 10−9 is more standard) | 1×10−8 | 1×10−8 |
| | homeologous exchange | 0 | 0 | 0 |
| | Num genes | Not applicable | 1250 | 1250 |
| Population | Na | 10^5-10^6 | 2x10^6 | 2x10^4 |
| | Nb | 10^4 | 10^4 | 100 |
| | burnin time | Not applicable | 4x10^8 | 4x10^6 |
| Seq. Evolution | WGD_gene_half_life | 31MY | 3e7 | 3e5 |
| | mutation_rate | 7.77 x 10-9 | 7.77x10-9 | 7.77x10-7 |
| Speciation | DIV time | 1.44 - 0.675 MYA | 600000 | 6000 |
| | WGD time | 0.35 - 0.610 MYA | 300000 | 3000 |

Num SSD genes simulated: 4371
Num paralogs in final genome: 5620
Q-scaling (reduce population & increase time-dependent rates) was used to decrease simulation time to under one hour. Q-scaling is discussed in the Supp. Methods.

| **Supp. Table S17. DemographiKS parameters for *Zea mays***<br>(a) Genes w/o HomEx=EMP_Mays_35_m09d23y2025_h14m59s18<br>(b) Genes w HomEx=EMP_Mays_36_m09d23y2025_h14m59s21 | | | | | |
|---|---|---|---|---|---|
| | | Est. from prior works | DGKS Q1 (theory) | DGKS Q100 35 | DGKS Q100 36 |
| chromosome | total_num_bases | 11.4M | 11.4M | 5.0M | 2.5M |
| | num_codons_in_a_gene | 1500 | 1500 | 1500 | 1500 |
| | recombination_rate | 7.3*10^-9 | 7.3*10^-9 | 7.3e−7 | 7.3e−7 |
| | homeologous exchange | Segmental. Varies across chr, 0-0.5 | 0,0.5 | 0.0000075 | 0.25 |
| Population | Na | 10^5 | 10^5 | 1e3 | 1e3 |
| | Nb | 1000 (bottleneck) max at 10^7 | 10^5 | 2e3 | 2e3 |
| | burnin time | Not applicable | 10^7 | 5e5 | 5e5 |
| Seq. Evolution | WGD_gene_half_life | 5-31 My | 3e7 | 3e5 | 3e5 |
| | mutation_rate | 2.17-3∗10−8 | 2.17e−8 | 2.17e−6 | 2.17e−6 |
| Speciation | DIV time | 11.9-20.5 mya | 11.9e6 | 11.9e4 | 11.9e4 |
| | WGD time | 5-12 Mya | 11.4e6 | 11.4e4 | 11.4e4 |
| Num SSD genes simulated: 13304<br>Num paralogs in final genome: 14967<br>Q-scaling (reduce population & increase time-dependent rates) was used to decrease simulation time to under two hours. Q-scaling is discussed in the Supp. Methods. | | | | | |

**Supp. Table S18. DemographiKS parameters for *Populus trichocarpa***
(a) Genes w/o HomEx=EMP_Pop_19_m10d01y2025_h17m31s43
(b) Genes w HomEx= EMP_Pop_18_m10d01y2025_h17m29s53

| | | Est. from prior works | DGKS Q1 (theory) | DGKS Q100 19 (used) | DGKS Q100 18 (used) |
|---|---|---|---|---|---|
| chromosome | total_num_bases | 13,000,000 | 13,000,000 | 1,500,000 | 1,500,000 |
| | num_codons_in_a_gen e | 1300 | 1300 | 1300 | 1300 |
| | recombination_rate | (4.2-7.8 )*10^-8 | 1e-8 | 1e-8 | 1e-8 |
| | homeologous exchange | Probably varies for diff chr, but low by now | 0-0.1 | 0 | 0.1 |
| Population | Na | ??? | | 10000 | 10000 |
| | Nb | 4000–6000 | 5000 | 2000 | 2000 |
| | burnin time | NA | 200000 | 200000 | 10000 |
| Seq. Evolution | WGD_gene_half_life | 3e7 | 3e7 | 3e5 | 3e5 |
| | mutation_rate | (4.2–5.2)×10^-8 | 4.2e-8 | 4.2e-6 | 4.2e-6 |
| Speciation | DIV time | ~4*10^6 | ~4.0e6 | 4.2e4 | 4.2e4 |
| | WGD time | ~4*10^6 | ~4.0e6 | 4.0e4 | 4.0e4 |

Num SSD genes simulated: 6918
Num paralogs in final genome: 9224
Q-scaling (reduce population & increase time-dependent rates) was used to decrease simulation time to under one hour. Q-scaling is discussed in the Supp. Methods.

**Supp. Table S19. DemographiKS parameters for *Saccharum spontaneum***
(a) Genes w/o HomEx=EMP_Sac_45_m09d19y2025_h10m27s28
(b) Genes w HomEx=EMP_Sac_46_m09d19y2025_h10m31s55

| | | Est. from prior works | DGKS Q1 (theory) | DGKS Q1000 (used) |
|---|---|---|---|---|
| chromosome | total_num_bases | chr varies from 100-44 MB | 2,000,000 | 2,000,000 |
| | num_codons_in_a_gene | 1000 (simplifying from Tiley) | 1000 | 1000 |
| | recombination_rate | 8.97e-10 ~1e-9 | 8.97e-10 ~1e-9 | 1e-8 |
| | homeologous exchange | Frequent | 0.5 | (a) 0.0<br>(b) 0.5 |
| Population | Na | > e^6? | 10*e^6 | 10000 |
| | Nb | 0.01-0.5e6 | 4*10^6 | 4000 |
| | burnin time | Not applicable | 500000 | 500000 |
| Seq. Evolution | WGD_gene_half_life | 30 *10^6 | 30 *10^6 | 3e4 |
| | mutation_rate | 3.8 × 10−9 (The complex polyploid genome architecture of sugarcane) | 3.8 × 10−9 | 3.8e-6 |
| Speciation | DIV time | 1.5 MY, sugar WGD, 70 MY ancient cereals WGD | 40*10^6 | 40000 |
| | WGD time | | 40*10^6 | 40000 |

Num SSD genes simulated: 13320
Num paralogs in final genome: 14652
Q-scaling (reduce population & increase time-dependent rates) was used to decrease simulation time to under two hours. Q-scaling is discussed in the Supp. Methods.

| Supp. Table S20. DemographiKS configurable parameters | | |
|---|---|---|
| **Parameter** | **Default Setting** | **Explanation** |
| Paths<br>> output_folder_root | >/home/DemographiKS_output | Destination for all output files |
| > pre_existing_trees_file | > False | Debugging option to use a pre-computed trees file. Saves time. |
| Chromosome:<br>> total_num_bases<br>> num_codons_in_a_gene | Chromosome:<br>> 2000<br>> 1000 | The total chromosome length in base pairs and the number of codons in a gene. This is used to set up SLiM. |
| > recombination_rate | > 1.258e-6 | The recombination rate units are per bp and per generation. Goes to SLiM. |
| > homoeologous_exchange_rate | > 0 | The homoeologous exchange<br>Rate between subgenomes. This is exposes the "Dij" parameter from<br>(Blischak et al. 2023). Goes to SLiM. |
| > max_num_paralogs_to_process | > False | Debugging option to only process the first few paralogs |
| Population:<br>> ancestral_ne (Na)<br>> bottleneck_ne (Nb) | Population:<br>> 100<br>> 20 | The ancestral diploid population size and Nb, the polyploid population size after WGD. |
| > burnin_time | > 2000 | As needed for to achieve steady state |
| Population:<br>> WGD_gene_half_life<br>> mutation_rate | Population:<br>> 31e-6<br>> 1.0e-5 | The WGD_gene_half_life units are per gene and per generation.The mutation rate units are per bp and per generation. |
| Mating:<br>> assortative_mating_coefficient | > 0 | This value can vary from 0 to 1, where 0 is panmixia and 1 is strong preferential mating for those with similar pedigree. |
| Migration:<br>> mig_start_gen | Migration:<br>> False | "mig_start_gen" is the generation at which to start migration. "Mig_stop_gen" is the |

| | | |
|---|---|---|
| > mig_stop_gen<br>> mig_rate | > False<br>> False | generation to stop migration. It's relative to the parental div time. “mig_rate” is the migration rate. Ie, 0.1 is 10% exchange of individuals. |
| Speciation:<br>> DIV_time_Ge<br>> WGD_time_Ge | Speciation:<br>> 4000<br>> 2000 | Time of parental divergence (False for autopolyploids) and time of whole genome duplication (in generations) |
| Randomization:<br>> SLiM_rep<br>> Msprime_random_seed<br>> DemographiKS_ random_seed | Randomization:<br>> 1<br>> 42<br>> 17 | Random seeds that are passed to each component of DemographiKS. These affect draws from random distributions used by the different components. Keep these numbers the same for total reproducibility. |
| Misc/Debugging:<br>> SimName<br>> KeepIntermediaryFiles<br>> StopAtStep<br>> AncestralGenomes ToSample | Misc/Debugging:<br>>allotetraploid_ bottleneck<br>> False<br>> 999<br>> [[1,5]] | “SimName” can be whatever you want. Setting “KeepIntermediaryFiles” to true will retain all the .fa files of your paralogs, and take up a lot of space. “StopAtStep” allows you to only run to a certain step (1,2,3,4,5,6). “AncestralGenomes ToSample” allows you specify which genome-pairs you want to sample, when building the ancestral coalescence. Ie, you might choose [[1,5],[3,4]] to check they both give the same distribution. |
